# Deglutarylation of GCDH by SIRT5 controls lysine metabolism in mice

**DOI:** 10.1101/2020.06.28.176677

**Authors:** Dhaval P. Bhatt, C. Allie Mills, Kristin A. Anderson, Bárbara J. Henriques, Tânia G. Lucas, Sara Francisco, Juan Liu, Olga R. Ilkayeva, Alexander E. Adams, Shreyas R. Kulkarni, Donald S. Backos, Paul A. Grimsrud, Cláudio M. Gomes, Matthew D. Hirschey

## Abstract

A wide range of protein acyl modifications has been identified on enzymes across various metabolic processes; however, the impact of these modifications remains poorly understood. Protein glutarylation is a recently identified modification that can be non-enzymatically driven by glutaryl-CoA. In mammalian systems, this unique metabolite is only produced in the lysine and tryptophan oxidative pathways. To better understand the biology of protein glutarylation, we studied the relationship between enzymes within the lysine/tryptophan catabolic pathways, protein glutarylation, and regulation by the deglutarylating enzyme Sirtuin 5 (SIRT5). Here, we identify glutarylation on the lysine oxidation pathway enzyme glutaryl-CoA dehydrogenase (GCDH). We show increased GCDH glutarylation when glutaryl-CoA production is stimulated by lysine catabolism. Our data reveal glutarylation of GCDH impacts its function, ultimately decreasing lysine oxidation. We then demonstrate the ability of SIRT5 to deglutarylate GCDH, restoring its enzymatic activity. Finally, metabolomic and bioinformatic analyses indicate an expanded role for SIRT5 in regulating amino acid metabolism. Together, these data support a model whereby a feedback loop exists within the lysine/tryptophan oxidation pathway, in which glutaryl-CoA is produced, in turn inhibiting GCDH function *via* glutaryl modification of GCDH lysine residues, and can be relieved by SIRT5 deacylation activity.

## INTRODUCTION

Protein post-translational modifications (PTM) are an evolutionarily conserved mechanism of cellular control across species (1–3). The discovery of several novel protein acyl modifications has expanded the spectrum of known PTMs in biology, including lysine propionylation, malonylation, butyrylation, beta-hydroxybutyrylation, succinylation, glutarylation, hydroxymethylglutarylation, methylglutarylation, methylglutaconylation, crotonylation, myristoylation, and palmitoylation [see (1, 2, 4) for reviews]. Large-scale proteomic studies have identified several biological targets of protein acylation, enhancing our understanding of the mechanisms and biological conditions leading to these acyl-modifications (4–6).

Protein acetylation and acylation in the nucleus and cytoplasm is primarily facilitated by acyl-transferase enzymes; however, under conditions of elevated pH such as in the mitochondrial environment, enzyme acylation can occur non-enzymatically. Thus far, in the absence of *bona fide* mitochondria acyl-transferases, the primary mechanism for mitochondrial protein acylation is generally considered to be non-enzymatic. Metabolite-based reactive carbon species (RACS) are often generated as metabolic intermediates, with a number of PTMs corresponding to their cognate acyl-CoA species.

While protein modification via RACS is emerging as nonenzymatic, acylation removal is enzymatically catalyzed by sirtuin protein deacylases. Sirtuins are a class of enzymes associated with stress response and aging(7, 8). The mitochondrial sirtuins, SIRT3, SIRT4 and SIRT5, have an expanding repertoire of deacylase activities; however, the biological roles of the mitochondrial sirtuins, and the acyl-modifications they regulate, remain unclear.

We previously showed that mice lacking SIRT5 have hyperglutarylated mitochondrial proteins (9), which play a key role in regulating the enzyme CPS1 in ammonia detoxification and the urea cycle (10). The only known source of glutarylation is glutaryl-CoA, a 5-carbon metabolite exclusively produced in the lysine (KEGG: hsa00310)/tryptophan (KEGG: map00380) catabolic pathways in mammalian systems (11). Furthermore, we previously identified glutaryl-CoA as a reactive carbon species (4), thus we predicted that SIRT5-mediated removal of protein glutarylation might control enzymes activity in the glutaryl-CoA metabolism pathway. Thus, we set out to test this hypothesis.

## RESULTS

Because of the emerging idea that proteins in the vicinity of reactive acyl-CoA’s are susceptible to non-enzymatic acylation of lysine residues, we explored protein glutarylation in the lysine/tryptophan degradation pathways. Within these pathways α-ketoadipate is converted to glutaryl-CoA by a protein-complex involving 2-oxoglutarate dehydrogenase (OGDH)/ dihydrolipoyllysine-residue succinyltransferase (DLST)/ dihydrolipoyl dehydrogenase (DLD). However, none of these protein-complex components are unique to the lysine/tryptophan degradation pathways, which might obfuscate our test of this hypothesis. Further downstream in this degradation pathway, glutaryl-CoA is converted into crotonyl-CoA by GCDH, which is exclusive to the lysine/tryptophan degradation pathways. Therefore, we selected GCDH as a putative target protein for further testing.

### GCDH is hyperglutarylated in the absence of SIRT5

To determine the acylation state of GCDH, we first overexpressed a C-terminal FLAG-tagged human GCDH protein (GCDH-FLAG) in HEK293T cells under basal cell culture conditions (high glucose DMEM, containing 4 mM glutamine, and 10% FBS). Overexpressed GCDH-FLAG was immuno-precipitated using a FLAG-M2 antibody-conjugated resin and probed for glutaryl-lysine using immuno-blot. We found that overexpressed GCDH-FLAG was glutarylated (Figure 1A). To confirm this finding, we performed a reverse immuno-precipitation experiment. Glutarylated proteins were immuno-precipitated using an anti-glutaryl-lysine antibody, and analyzed using immuno-blot analysis with anti-FLAG antibody. We found a distinct band detecting the FLAG-tag on overexpressed GCDH-FLAG at 43 kDa (Figure 1A), the molecular weight of mitochondrial GCDH. These data provide evidence that GCDH is acylated.

**Figure 1.**
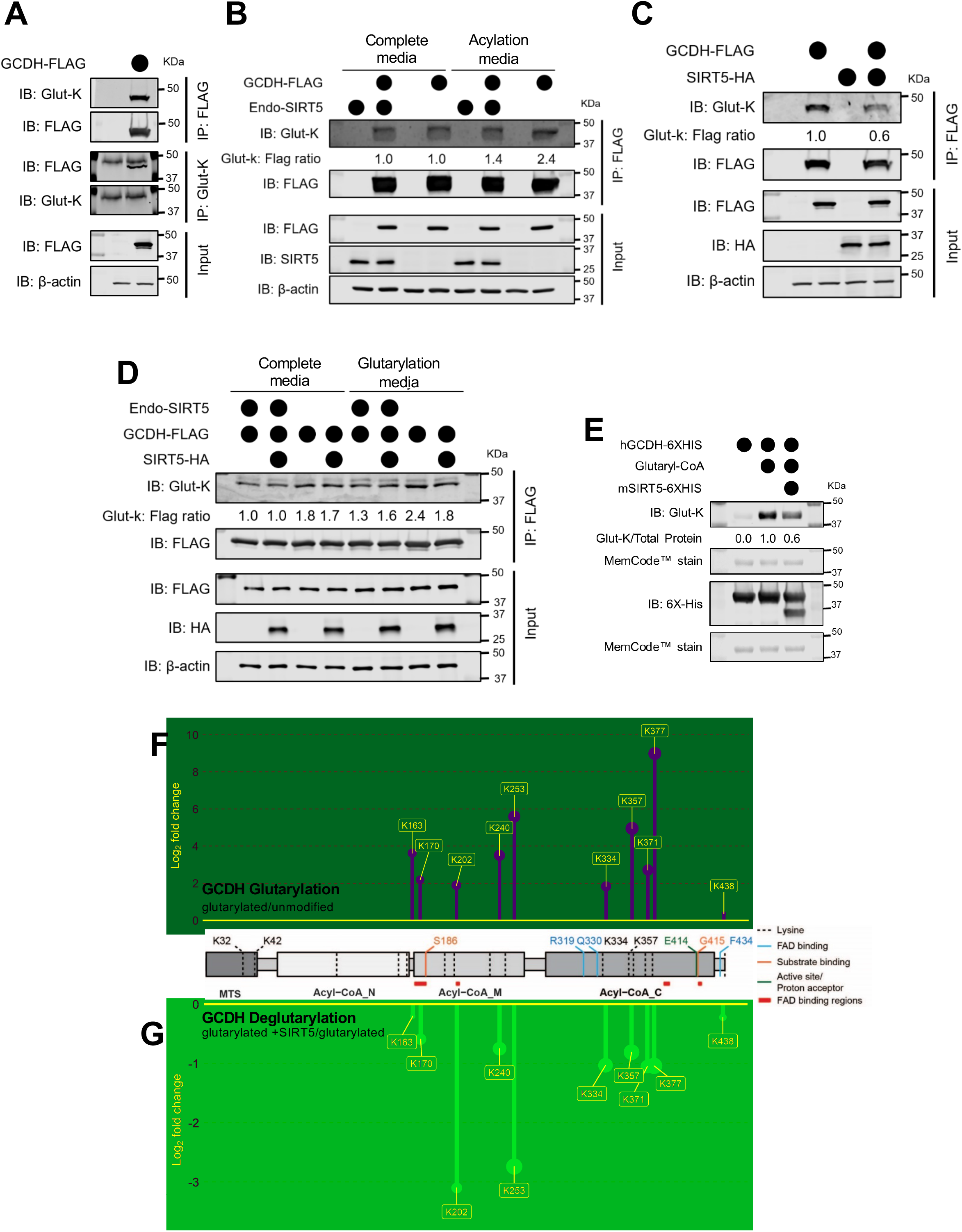
GCDH is de-acylated by Sirt5, restoring its enzymatic activity. **(A)** Immunoblot for immuno-purified GCDH-FLAG by Flag-M2 resin or glutaryl-proteins by anti-glutaryl-lysine antibody (Glut-K) from HEK293T cells grown in complete media. Blots representative of at least three independent experiments. **(B)** Immunoblot for immuno-purified GCDH-FLAG by Flag-M2 resin from SIRT5 crWT or crKO cells grown in complete media or acylation media (DMEM - glucose, - glutamine, - pyruvate, - 10% FBS). Blots are representative of at least three independent experiments and quantitative values of glutaryl-lysine intensity normalized to FLAG intensity expressed as ratios relative to control (overexpressed GCDH-FLAG in SIRT5 crWT cells grown in complete media, lane 2). **(C)** Immunoblot of immuno-purified GCDH-FLAG by Flag-M2 resin from HEK293T cells grown in complete media with or without co-expressed SIRT5-HA. Blots representative of at least three independent experiments and quantitative values of glutaryl-lysine intensity normalized to FLAG intensity expressed as rations relative to control (overexpressed GCDH-FLAG, lane 2) **(D)** Immunoblot for immuno-purified GCDH-FLAG by Flag-M2 resin from SIRT5 crWT or crKO cells grown in complete media or glutarylation media (EBSS +5mM glucose, +50mM HEPES, +0.8mM Lysine) with or without co-expressed SIRT5-HA. Blots are representative of at least three independent experiments and quantitative values of glutaryl-lysine intensity normalized to FLAG intensity expressed as ratios relative to control (overexpressed GCDH-FLAG in SIRT5 crWT cells grown in complete media, lane 1) **(E)** Immunoblot of chemically modified GCDH (hGCDH-6xHIS) using glutaryl-CoA with or without incubation with recombinant SIRT5 (mSIRT5-6xHIS). Quantitative values of glutaryl-lysine intensity normalized to total protein are expressed as ratios relative to control (glutaryl-modified recombinant GCDH, lane 2) **(F)** Enzymatic activity of unmodified (black), glutarylated (dark purple) and deglutarylated (light purple) GCDH determined by using PMS/DCPIP as artificial electron acceptors and Glutaryl-CoA as electron donor (mean ± SD, n=6, 6, and 5 for unmodified, modified and deacylated GCDH respectively; * = p-value ≤ 0.05). **(GH)** Glutaryl- lysine sites identified by label-free quantitative LC-MS/MS on recombinant GCDH mapped to full-length human GCDH protein schematic with domain and residues important for cofactor, substrate, or tetramer interactions. Panel G shows glutaryl-lysine sites in chemically glutarylated GCDH sample expressed as ratio over unmodified GCDH (Glutarylation: glutarylated GCDH/unmodified). Pael H shows glutaryl-lysine sites on chemically glutarylated GCDH altered by SIRT5 incubation as expressed as a ration over glutarylated sample(Deglutarylation: glutarylated GCDH + SIRT5/glutarylated GCDH), Plots representing Log_2_ fold change (y-axis) and p-value (larger circle size = more significant p-value).

Next, we determined if SIRT5 influenced GCDH glutarylation levels. Of all the known sirtuin activities, SIRT5 is the only one known to possess strong deglutarylase activity (9), in addition to its desuccinylase and demalonylase activities (12–14). To test how SIRT5 influenced acylation of GCDH, we generated a SIRT5KO cell line using the CRISPR-Cas9 system in HEK293T cells (SIRT5 crKO). Five different crRNA’s targeting different regions upstream of the catalytic histidine in the exon 4 or 5 of the four known coding transcripts of *SIRT5* generated 75 monoclonal colonies. The control cell line (SIRT5 crWT) was processed through the same CRISPR-Cas9 workflow like the SIRT5 crKO cells except the *SIRT5* crRNA was omitted. We screened each colony for SIRT5 protein levels using immunoblot and found one colony (cr guide E, Table S1 with complete SIRT5 depletion (Supporting Figures 1A-C). Surprisingly, during screening, we noticed some colonies showing ~33% to 66% reduction of SIRT5 protein. Upon analyzing HEK293T karyotyping and copy number analysis (15), it became clear that these hypo-triploid cells have three copies of the human *SIRT5* gene. Using immunoblots, we monitored the SIRT5 crKO clone for SIRT5 protein levels over several passages and checked for changes in protein acylation. While no SIRT5 protein was detected in the SIRT5 crKO cell line for at least 15 passages (data not shown), we found modest increases in global protein glutarylation, succinylation, malonylation and acetylation relative to the control SIRT5 crWT line (passage 10, Supporting Figure 1D-K). Thus, this SIRT5 crKO cell line appeared to be a suitable model to study the role of SIRT5.

Next, we overexpressed GCDH-FLAG in SIRT5 crKO cells, predicting the absence of SIRT5 would lead to an increase in glutarylation of GCDH. However, we did not find any significant change in the glutarylation state of overexpressed GCDH-FLAG in SIRT5 crKO cells under basal cell culture conditions (Figure 1B). Since cells were grown in complete media with abundant nutrients, we hypothesized the cells might not be metabolizing lysine/tryptophan; the only known pathways that generate glutaryl-CoA. Therefore, we reasoned that under basal cell growth conditions, glutaryl-CoA levels may not be produced in quantities high enough to induce hyper-glutarylation. We repeated the same experiment in cell culture media designed to drive catabolism of amino acids and ultimately acylation, here termed acylation media (DMEM, - glucose, - glutamine,- pyruvate, - serum), and assessed the glutarylation-state of overexpressed GCDH-FLAG. Under these media conditions, the only oxidative fuels available to these cells were amino acids (excluding glutamine), which drive acyl-CoA formation and result in increased acylation. We found that increasing amino acid oxidation in this media increased glutarylation of GCDH-FLAG in both SIRT5 crWT and crKO cells relative to cells grown in complete media, however to a much higher degree in the SIRT5 crKO cells (2.4-fold) compared to the SIRT5 crWT cells (1.4-fold; Figure 1C). Therefore, these data suggest that limiting the availability of non-amino acid fuel sources increases glutarylation of GCDH in SIRT5 crWT cells, which further increases in absence of the protein deacylase SIRT5.

To determine the specificity of this model to glutaryl-CoA, we tested whether succinylation, another modification targeted by SIRT5, modifies GCDH under conditions similar to glutarylation. We investigated whether absence of SIRT5 altered the succinylation-state of overexpressed GCDH-FLAG using identical conditions tested for glutarylation in SIRT5 crWT and SIRT5 crKO with complete and acylation media. Unlike glutaryl-GCDH, we found that under basal conditions GCDH-FLAG was hypersuccinylated in SIRT5 crKO cells (Supporting Figure 2A). Our previous work has shown that succinyl-CoA is more reactive than glutaryl-CoA, as it has greater ability (approximately 10-fold) to form a cyclic anhydride. Supporting these data, succinyl-CoA modified proteins show greater molecular weight shifts(4) compared to glutaryl-CoA or any other 4-5 carbon dicarboxylic acid. Thus, basal hyper-succinylation of GCDH-FLAG could be due to elevated reactivity of succinyl-CoA and/or its higher cellular abundance. Moreover, similar to glutaryl-GCDH, starvation further increased succinylation of GCDH-FLAG in both SIRT5 crWT and SIRT5 crKO cells, again to a greater extent in SIRT5 crKO cells (3.5-fold) than SIRT5 crWT cells (1.7 fold) (Supporting Figure 2A).

The acylation medium used in these experiments had normal levels of all amino acids except glutamine. Under these conditions, in absence of the preferred fuel sources (glucose, glutamine, and pyruvate), cells are forced to metabolize amino acids for energy production. Strikingly, both succinyl-CoA-generating (derived from valine and isoleucine) and glutaryl-CoA-generating (derived from lysine and tryptophan) amino acids were available to the cells and could potentially explain the increase in succinyl- and glutaryl-modifications of GCDH during starvation.

### SIRT5 deglutarylates GCDH

To further test the relationship between GCDH and SIRT5-mediated deglutarylation, we co-overexpressed GCDH-FLAG and a C-terminal HA-tagged human SIRT5 in HEK293T cells. GCDH-FLAG, which was affinity purified from cells overexpressing SIRT5-HA showed significantly less (0.6-fold) glutarylation compared to that from cells expressing endogenous levels of SIRT5 (Figure 1C). Interestingly, while testing the ability of overexpressed SIRT5 to desuccinylate GCDH-FLAG, we found that unlike glutaryl-GCDH, co-expression with SIRT5-HA did not reduce the succinylation state of GCDH-FLAG (Supporting Figure 2D). These data suggest that SIRT5 targets glutaryl-GCDH for deglutarylation, but does not appear to alter the succinylation state of GCDH as measured by immunoblotting.

To interrogate the specificity of SIRT5 towards glutaryl-GCDH, we performed GCDH-FLAG and SIRT5-HA overexpression in SIRT5 crWT and crKO cells in complete and glutarylation promoting conditions. To do so, we created a glutarylation media composed of Earle’s Balanced Salt Solution (EBSS, with 5mM glucose) supplemented with 50 mM HEPES and 0.8 mM lysine (as a source of glutaryl-CoA). Our rationale was to grow cells in a medium with minimal fuel sources to drive lysine oxidation and generate substrate for glutarylation (glutaryl-CoA) rather than in media that generates precursors for both glutarylation and succinylation (succinyl-CoA), as occurs in the above-mentioned acylation-promoting medium. Interestingly, we found that SIRT5-HA was only able to deglutarylate GCDH-FLAG under these nutrient deprivation conditions in the absence of endogenous SIRT5 (SIRT5 crKO; Figure 1D). As expected, the increase in succinylation of GCDH in glutarylation media was not as pronounced as in acylation promoting media (Supporting Figure 2A vs B). Absence of succinyl-CoA generating amino acids in the glutarylation media may explain the differences in succinylation of GCDH-FLAG compared to the marked increase (~3.5 fold) observed in acylation media (Supporting Figure 2A vs B). However, consistent with the SIRT5 and GCDH co-expression in HEK93T cells, overexpressed SIRT5 modestly desuccinylated GCDH-FLAG (Supporting Figure 2B). Overall, these data support a role for SIRT5 in deglutarylating GCDH, but not desuccinylating the protein under nutrient deprivation conditions.

### Glutaryltion impact on GCDH conformation

To identify the sites of acylation on GCDH, we turned to a well-established mass spectrometry-based proteomic approach using chemically modified and unmodified recombinant GCDH. GCDH protein was separated by SDS-PAGE, in-gel digested using trypsin, and glutaryl-or succinyl-GCDH peptide relative abundance was measured and normalized to GCDH proteins measured from unmodified peptides. Using this strategy, we obtained 99% sequence coverage of GCDH protein and all lysine residues corresponding to the cleaved mitochondrial protein were identified with acyl modifications (Table S2). Recombinant GCDH protein used for this analysis lacked the mitochondrial localization sequence, however all residue locations referred in the figures and text correspond to the full-length human GCDH protein. Overall, we found 10 glutaryl-lysine (Figure 1G), and 11 succinyl-lysine (Supporting Figure 2G) sites on GCDH. Nine lysine residues were significantly glutarylated, with > 1.8 Log_2_ fold-change above unmodified GCDH (p ≤ 0.05). Of those glutarylated sites, six were significantly deglutarylated (p ≤ 0.05; Figure 1H), with Lysine 253 having the greatest significant fold-change compared to glutarylated GCDH. These data further support regulation of GCDH glutarylation by SIRT5.

To gain insight into how glutarylation influences GCDH, we performed molecular modeling combined with a battery of biophysical measurements. GCDH functions as a homo-tetramer with substrate (glutaryl-CoA, light gray spheres) and co-factor (FAD, black spheres) binding pockets (Supporting Figure 3D-E). Modeling of acyl sites on the crystal structure of human GCDH (PDB ID: 1SIR) showed that several of these sites are important for tetramer stability (K253, K335, K361, and K377), substrate binding (K111, K163, K170, K210, K357, K361, K371 and K438) and cofactor binding (K163, K170, K210, K253, K335, and K371). Notably, all heavily glutarylated lysine residues (163, 170, 253, 357, 371, and 377) except K202 and K240 have a strong potential to influence FAD/glutaryl-CoA binding and tetramer formation.

K163 and 253 both form important bonds involved in GCDH co-factor binding, and modification of these residues could impact such binding. Furthermore, K253, along with K377, resides in the protein monomer interaction interface within GCDH, potentially influencing tetramer stability (Supporting Figure 3D-E). Additionally, K163, 170 and 240 are all involved in salt bridges that likely important to overall protein structure. While a single modification at one of these locations may have limited effect on protein structure, modification of all three simultaneously could reduce the overall stability of the protein structure and could indirectly affect substrate binding as this region is on the opposite side of the protein from the binding pocket (Supporting Figure 3E).

To test the structural impact of acylation on GCDH enzyme as suggested by molecular modeling, we performed extensive conformational and stability studies in purified heterologous expressed human GCDH (see materials and methods for details) using different spectroscopic techniques which provide information on different levels of protein structure. To assess the effect on GCDH’s secondary structure, we used circular dichroism (CD) in the far-UV region. We found that glutarylation results in a small decrease in ellipticity suggesting a slight change in secondary structure content (Supporting Figure 3G). Furthermore, when monitoring thermal unfolding following changes in secondary structure (ellipticity at 222nm), we observed a slight decrease (Δ T_m_ = −3 °C) in the thermal stability of glutarylated GCDH compared to that of unmodified GCDH, which has a melting temperature (T_m_) of 55 ±1 °C (Supporting Figure 3H). Parallel studies using succinylated GCDH showed a similar profile: a slight change in secondary structure content and decreased stability for modified protein (Supporting Figure 3I-J). We also monitored modified GCDH using fluorescence tryptophan emission to monitor tertiary structure and FAD emission to assess the cofactor environment. These studies showed no relevant alteration on protein conformation, as judged by spectra at 25°C, or on protein thermal stability, as evaluated by thermal denaturation curves, irrespective of modifications (data not shown).

Furthermore, glutarylation abrogates the +1 charge present on lysine side-chains, leaving a −1 charged glutaryl-modification, which is sterically larger than unmodified lysine. These changes in the steric space, lysine charge, or both, could disrupt interactions between protein subunits or protein-protein interactions. Since functional GCDH occurs as a homo-tetramer, we investigated whether altered steric and/or charged states could influence the native tetramer-complex of GCDH. To do so, we isolated mitochondria from SIRT5WT and SIRT5KO mouse livers and performed blue-native polyacrylamide gel electrophoresis (BN-PAGE) to monitor the GCDH tetramer. Identity of correct GCDH tetramer band was confirmed by using GCDHWT and GCDHKO mouse liver mitochondrial lysates in parallel (Supporting Figure 3C). In the fed state, we did not observe any significant change in the molecular weight of the GCDH tetramer between the two genotypes (Supporting Figure 3C, quantified 3F). Indeed, this agrees with size exclusion chromatography assays performed with purified GCDH (unmodified and modified) which showed no difference in protein quaternary structure (data not shown). Since our previous analysis showed that acylation of GCDH increases during starvation, we isolated liver mitochondria from 24 h fasted SIRT5WT and KO mice and repeated the BN-PAGE analysis. We observed no major differences between GCDH tetramer in SIRT5KO and SIRT5WT mouse liver mitochondria (data not shown). Therefore, although structural modeling suggests that tetramer formation or cofactor biding could be influences by acylation, *in vitro* and *ex vivo* analysis showed that glutarylation has no major impact on the native GCDH folding.

### Lack of SIRT5 impairs glutaryl-CoA and lysine oxidation

As described above, GCDH is an enzyme involved in the lysine and tryptophan oxidation pathways. Lysine is one of the most abundant amino acids found in the body that can be used for ketogenesis during periods of starvation (16). During starvation, lysine is metabolized to acetyl-CoA and CO_2_ (Figure 2A), which can be used to fuel the TCA cycle or used for other metabolic processes. Thus, we tested the impact of SIRT5 loss on lysine catabolism. First, we investigated the significance of hyperacylated GCDH on lysine oxidation. To do so, we modified an *in vitro* leucine oxidation assay (17) for measuring lysine metabolism using U-[^14^C]-labeled lysine. SIRT5 crWT and crKO cells were incubated in glutarylation medium (EBSS, +5mM glucose, + 50 mM HEPES, 4μCi/mL U-[^14^C]-lysine, 0.8 mM ^12^C-lysine) containing radiolabeled and unlabeled lysine for a period of 3 hours, followed by quantification of released [^14^C]-CO_2_. These measurements showed that SIRT5 crKO cells had significantly reduced oxidation of lysine (by 40%) as compared to the SIRT5 crWT cells (Figure 2B). These data support the notion that hyperacylation of GCDH can functionally impair flux through this pathway.

**Figure 2.**
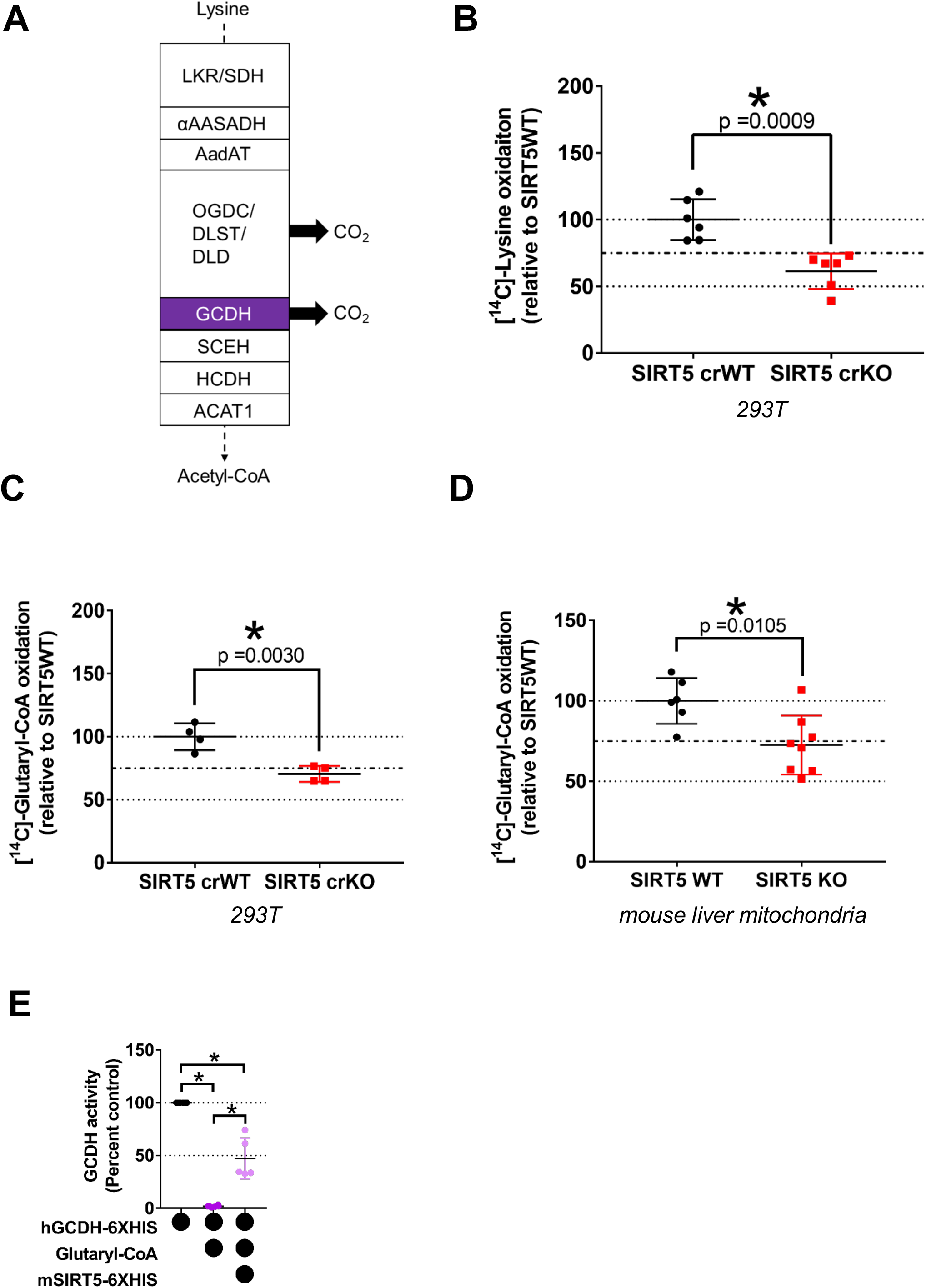
SIRT5 influences lysine oxidation. **(A)** Schematic of lysine oxidation enzymes with steps that release carbon dioxide (CO_2_). **(B)** Oxidation of [U-^14^C]-lysine to [^14^C]-CO_2_ in 293T SIRT5 crWT and crKO cells. Data representative of three independent experiments (mean ± SD, n = 6; **\***= p-value ≤ 0.05). **(C)** Oxidation of [1, 5-^14^C]-glutaryl-CoA to [^14^C]-CO_2_ in mitochondria isolated from 3h starved 293T SIRT5 crWT and crKO cells. Data representative of single experiment (mean ± SD, n = 4; **\***= p-value ≤ 0.05). **(D)** Oxidation of [1, 5-^14^C]-glutaryl-CoA to [^14^C]-CO_2_ in mitochondria isolated from SIRT5WT and SIRT5KO mouse liver. Data representative of single experiment (mean ± SD, n = 6, 8 for SIRT5WT and SIRT5KO respectively; **\***=p-value ≤ 0.05).

Since lysine oxidation has two discrete steps that generate CO_2_ (Figure 2A), either at the dehydrogenase complex described above or at GCDH, we wanted to determine if impaired lysine oxidation observed in SIRT5 crKO cells results from reduced GCDH activity directly. For this, we modified the above described lysine oxidation assay in a way that allowed us to directly quantify the ability of GCDH to convert glutaryl-CoA into crotonyl-CoA and CO_2_. Since glutaryl-CoA cannot enter living cells, we isolated mitochondria from the SIRT5 crWT and crKO cells that were treated glutarylation media, in a manner similar to lysine oxidation above. Mitochondria were permeabilized and incubated with [1, 5-^14^C]-glutaryl-CoA in presence of FAD, the co-factor essential for GCDH activity. After 1 hour of incubation, we found that SIRT5 crKO cell mitochondria had significantly reduced ability to release radiolabeled CO_2_ (~25%; Figure 2C). We further confirmed these observations *ex vivo*, by using permeabilized liver mitochondria from 24-hour fasted SIRT5WT and SIRT5KO mice and observed a similar reduction in GCDH activity (~25%; Figure 2D); GCDHKO mouse liver lysate served as a negative control for this assay (data not shown).

To better elucidate on acylation effect on enzyme function, and taking advantage of having a pure human GCDH protein model, we also measured the specific activity of the enzyme. For this *in vitro* assay, we determined the ability of acylated- and deacylated-recombinant GCDH to transfer electrons from glutaryl-CoA oxidation to either an artificial electron acceptor (phenazine methosulfate, PMS) or to its endogenous electron acceptor (electron transfer flavoprotein, ETF). Interestingly, we found that chemically glutarylated GCDH had almost undetectable activity using the artificial electron acceptor PMS and about 20% residual activity with ETF, when compared to unmodified GCDH (Figure 2E and S3A). These results thus indicate that acylation potently compromises GCDH function. Incubation of modified GCDH with recombinant SIRT5 partially restored (~50%) GCDH function (Figure 2E and S3A) in both conditions, in agreement with previous findings that SIRT5 is able to deglutarylate sites that are important for GCDH function. Consistently, succinylated GCDH also reduced GCDH activity to about 10% of unmodified GCDH, and incubation with SIRT5 was able to restore it to about 30% (Figure S2F and S3B).

Overall, these data suggest that impaired GCDH activity could contribute to the reduced ability of SIRT5 crKO cells to metabolize lysine, and more specifically, glutaryl-CoA. Additionally, although SIRT5 can target both glutaryl- and succinyl-lysine on GCDH, SIRT5 has a greater impact on deglutarylation of GCDH, and can modulate its function.

### Amino acid levels are increased in the absence of SIRT5

Because our previous results showed a decrease in lysine oxidation capacity in the absence of SIRT5, we asked whether metabolism of other amino acids was also altered in the absence of SIRT5. To test this, we performed both targeted and non-targeted metabolomics. When we measured total levels of amino acids in SIRT5 crWT and crKO cells, we found that the majority of amino acids exhibited increased levels in the absence of SIRT5 (Figure 3A), suggesting less consumption (oxidation or protein synthesis). We hypothesized that levels of lysine intermediates, along with those of other elevated amino acids, would be altered by SIRT5 ablation. To measure lysine and its intermediates, we performed non-targeted metabolomic analyses on SIRT5 crWT and crKO cells in the fed (complete DMEM), fasted (1h EBSS only) and refed (fasted 1h EBSS, refed 1h complete DMEM) states. We found significant changes in some lysine catabolic intermediates in SIRT5 crKO fed and fasted conditions as compared to the SIRT5 crWT cells (Figure 3B-D, pink). Importantly, these changes were not limited to lysine metabolism; we also observed significant changes in other amino acid metabolic intermediates (Figure 3B-D, pink). These data suggest the SIRT5 crKO phenotype goes beyond lysine catabolism, implying a broader role for SIRT5 in controlling amino acid metabolism.

**Figure 3.**
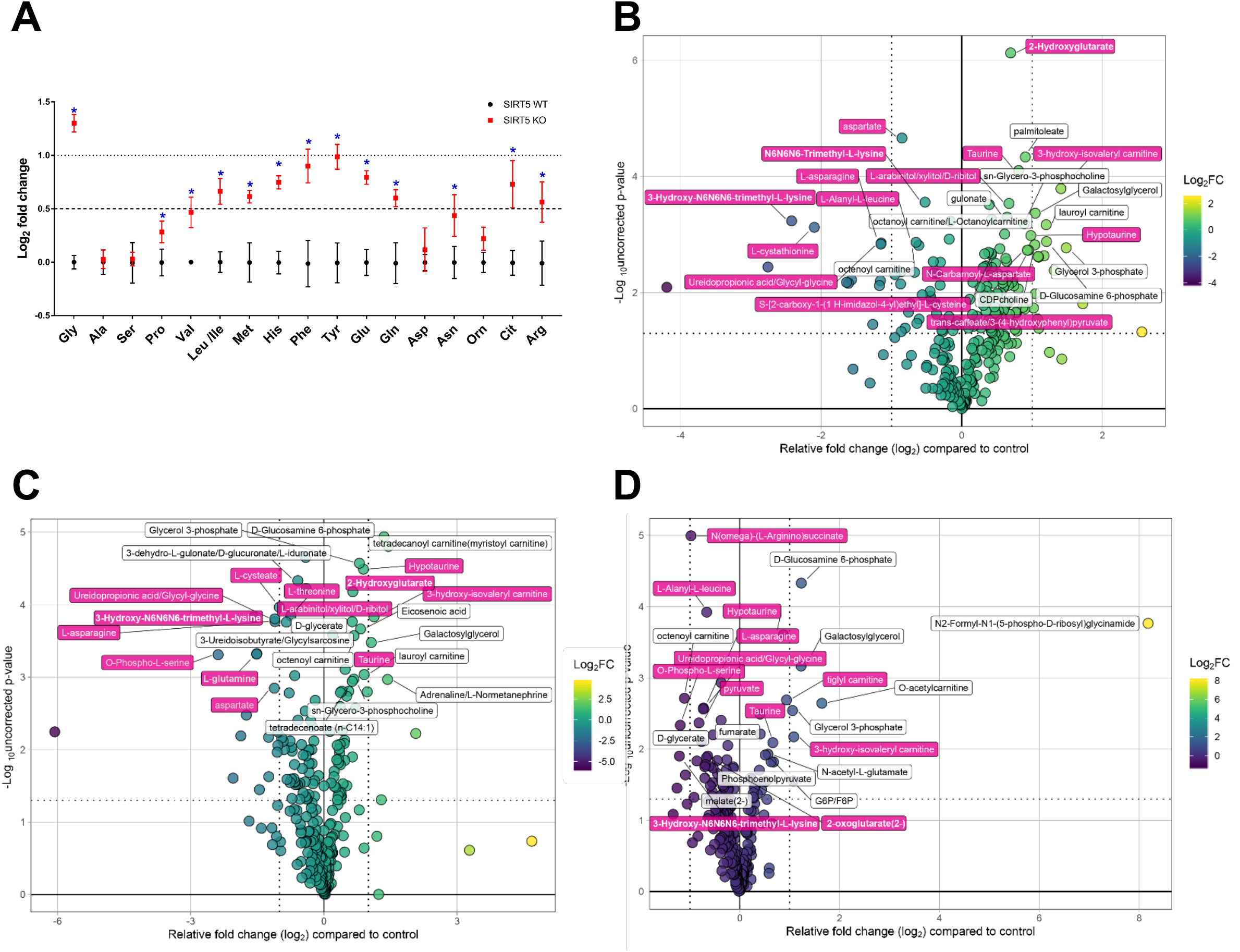
Amino acid levels are increased in the absence of Sirt5. **(A)** Targeted metabolomics in 293T SIRT5 CRISPR cells reveal increased levels in the majority of amino acids in crKO compared to crWT. **(B-D)** Non-targeted metabolomics in 293T SIRT5 crWT and crKO cells that have been fed (B), fasted (C) or refed (D). Plots showing relative fold change of 293T SIRT5 crKO compared to crWT reveal several amino acid metabolic intermediates (pink) are among the metabolites with the largest change between genotypes.

### Hepatic *SIRT5* expression correlates with amino acid metabolic genes

To further test the role of SIRT5 in amino acid metabolism, we performed gene expression correlation analysis using a human liver gene set. We identified a publicly available human liver gene set (NCBI GEO id 14520) measuring possible biomarkers for hepatocellular carcinoma. We filtered this dataset to only include non-tumor data, and performed a gene correlation analysis. We then searched for genes with the highest correlation to *SIRT5.* We found enrichment of genes involved in amino acid metabolism that positively correlated to *SIRT5* gene expression (Figure 4, red bars), meaning when *SIRT5* expression is high in human liver, expression of these genes is also high. When we looked at genes with a negative correlation to *SIRT5* expression, meaning when *SIRT5* expression is high these genes are low, we did not find enrichment of any particular processes. Furthermore, we preformed the correlation analyses for the other mitochondrial sirtuins, *SIRT3* and *SIRT4*, and did not find enrichment for amino acid metabolic processes, supporting a specific role for SIRT5 in amino acid metabolism (Supporting Figure 4).

**Figure 4.**
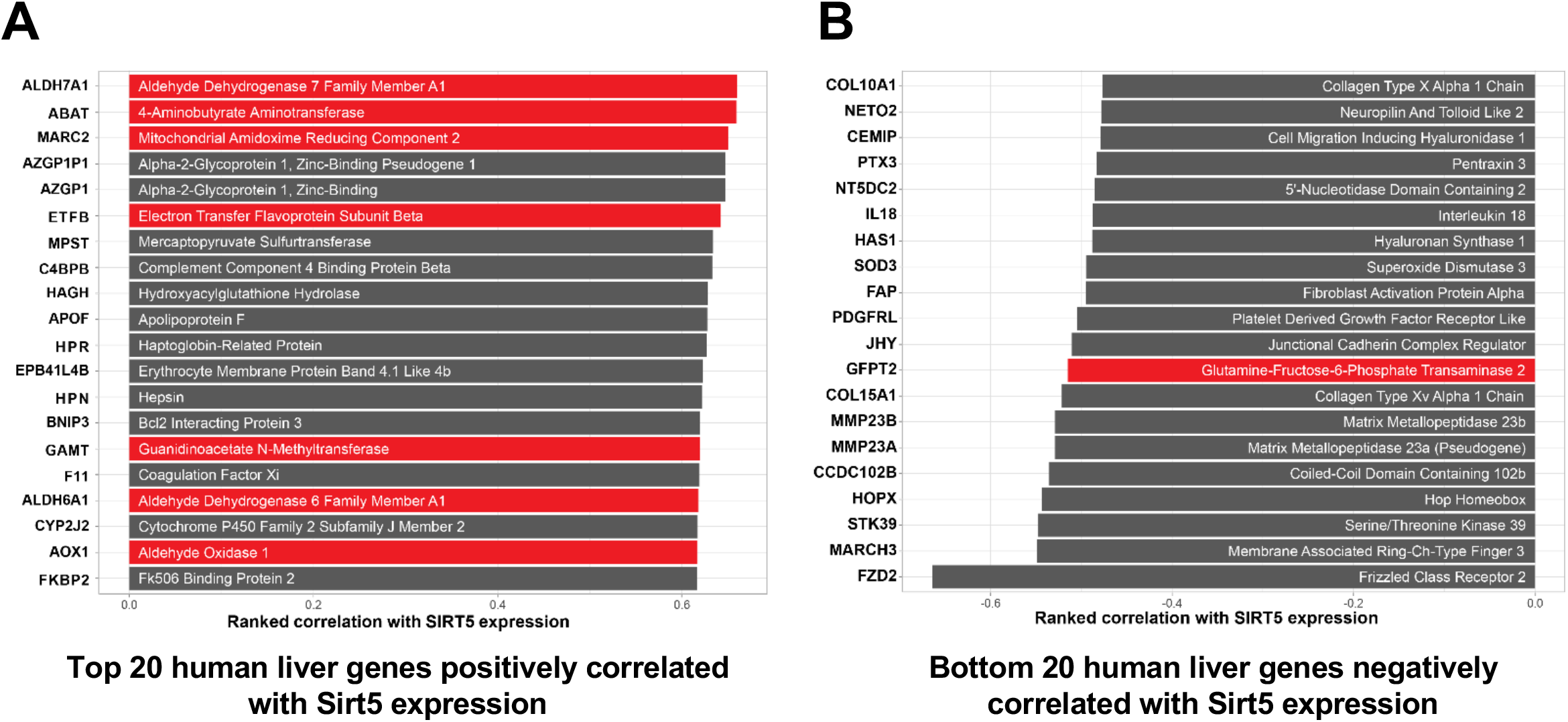
SIRT5 expression correlates with AA metabolism. **(A-B)** Human liver Sirt5 5 liver co-expression analysis. Gene expressions that positively (A) or negatively (B) correlate to Sirt5 gene expression. Genes highlighted in red have known roles in amino acid metabolism.

## DISCUSSION

Sirtuin 5 (SIRT5) is a mitochondrial and cytoplasmic NAD^+^-dependent protein deacylase that functions to remove a wide-range of protein modifications, including malonyl-, succinyl-, and glutaryl-lysine. While SIRT5 has been shown to deglutarylate CPS1, little is known about protein glutarylation and how SIRT5 functions to regulate this recently discovered modification. In this study, we found the glutaryl-CoA handling enzyme, GCDH, is glutarylated, and glutarylation increases under conditions promoting lysine/tryptophan metabolism (Figure 1). Furthermore, we demonstrated glutarylation of GCDH decreases enzymatic activity, which is partially relieved by de-glutarylation via SIRT5 (Figure 1F). These observations suggest a regulatory feedback loop within the lysine/tryptophan degradation pathways, in which increased flux through the pathway results in deactivation of the pathways via glutarylation of GCDH. Conversely, this inhibition of GCDH is relieved *via* deacylation by SIRT5,

Beyond identifying a novel substrate for SIRT5, we found a role for SIRT5 in controlling lysine metabolic flux. Additionally, these data indicate a potential role for SIRT5 in amino acid metabolism beyond lysine/tryptophan (Figures 3 and 4). When integrated with previous findings of known SIRT5 targets, we propose that SIRT5 could play a broader role in regulating feedback loops within amino acid metabolic pathways. For example, catabolism of branched-chain amino acids results in increased succinyl-CoA production, which in turn could increase protein succinylation. It is possible the enzymes within the amino acid catabolic pathways are among those succinylated proteins, and modification may impact their enzymatic functions. SIRT5 is a potent desuccinylase and could abrogate protein succinylation resulting from amino acid catabolism.

We previously explored the mechanism(s) underlying this non-catalytic nature of protein acylation and reported that a specific class of 4-5 carbon dicarboxylic acyl-CoA’s possess unusual reactivity compared to mono-carboxyl or shorter/longer dicarboxylic acyl-CoA species (4). We found that this enhanced reactivity results from an intrinsic property of these 4-5 carbon dicarboxylic acyl-CoA’s to undergo intra-molecular catalysis and form highly reactive anhydride intermediates. The pattern of protein acylation suggested that proteins/enzymes in pathways that generate the reactive acyl-CoA species become uniquely acylated by them. While it remains unknown whether enzymes that directly handle these acyl-CoA species undergo self-acylation and if this in turn regulates the function of the enzyme/pathway involved, the findings presented in this study support this model. Furthermore, the concept that proteins in the vicinity of acyl-CoA’s are susceptible to non-enzymatic acylation of lysine residues was first shown by methylcrotonoyl-CoA carboxylase (MCCC), an enzyme in the leucine oxidation pathway, becoming acylated by multiple acyl-CoA’s including 3-hydroxy-3-methylglutaryl-CoA (HMG-CoA), 3-methylglutaconyl-CoA (MGc-CoA), and 3-methylglutaryl-CoA (MG-CoA), all generated within leucine oxidation or ketogenesis pathways (17).

While we are far from a complete understanding of SIRT5, how RACS behave in biological systems, and the interaction between the two, the present study sheds some light on this unexpected regulatory mechanism in mitochondria. Interestingly, the role of SIRT5 in the cytoplasm or in the regulation of predominantly cytoplasmic modifications like malonylation remain enigmatic. Many studies have demonstrated that a pool of SIRT5 is found in the cytoplasm, however, we don’t yet know the function of cytoplasmic SIRT5. Moreover, we don’t know whether the stoichiometry of these acyl-CoA’s and the resulting modifications is enough to cause meaningful functional consequences. Additional studies will be needed to further understand the importance of SIRT5s deacylating activity, and to understand the widespread impact this system has on nutrient metabolism and cellular homeostasis.

## EXPERIMENTAL PROCEDURES

### Animals

Between 2-4 animals per cage were housed together with a 12 h light-dark cycle and received ad libitum PicoLab rodent diet 20 (#5053, LabDiet, St. Louis, MO) and water. All animal procedures were performed in accordance with the international guidelines from the Association for the Assessment and Accreditation of Laboratory Animal Care and approved by the Duke University Institutional Animal Care and Use Committee.

The SIRT5KO mice (B6;129-Sirt5tm1Fwa/J, stock #012757) were obtained from Jackson Laboratory (Bar Harbor, ME,) and were backcrossed for 10 generations with C57BL/6J mice (stock #000664, Jackson Laboratory, Bar Harbor, ME). Later, the resulting mice were backcrossed with the wild-type C57/BL6NJ (stock # 005304), strain from the Jackson Laboratory (Bar Harbor, ME) to obtain mice with a pure wild-type NNT background. The resulting mice heterozygous for SIRT5 but homozygous for wild type NNT were used as breeders to generate the SIRT5WT and SIRT5KO mice used in this study. For genotyping primers SIRT5WT forward (5’-AGG AGG TGG CAA AGG TCT TGC-3’), SIRT5KO forward (5’-TCA TTC TCA GTA TTG TTT TGC C-3’) and a common SIRT5WT/KO reverse (5’-CTG AGG TAG AGT CTC TCA TTG-3’) amplifying a 575 bp WT-SIRT5 band and/or a 350 bp KO-SIRT5 band were used to identify SIRT5WT and SIRT5KO genotypes.

The GCDHKO mouse strain (B6.129S4-Gcdhtm1Dmk/Mmnc, # 034368-UNC) was obtained from the Mutant Mouse Regional Resource Center (MMRRC, a NIH funded strain repository #8U42OD010924-13) that was donated to the MMRRC by David M. Koeller, MD, Oregon Health & Science University. These GCDHKO mice had a mixed C57/BL6J and C57/BL6NJ background and breeders heterozygous for the GCDH gene were generated by backcrossing the GCDHKO mice with the wild-type C57/BL6NJ strain (stock # 005304, Jackson Laboratory, Bar Harbor, ME). By sequential genotyping and selection of mice homozygous for WT-NNT and heterozygous for GCDH were bred to obtain the GCDHWT and GCDHKO mice used in this study. For genotyping, primer pairs GCDHWT forward (5’-CTT CCG TAA CTA CTG CCA GGA GCG G-3’) and GCDHWT reverse (5’-AGC TCT CGG GTC AGG AGC CCA TAG G-3’) and, GCDHKO forward (5’-TTA GGC CTA GTG TGC TGG TCC CGG A-3’) and GCDHKO reverse (5’-TCT GGT GCC GGA AAC CAG GCA AAG C-3’) were used to detect either a 565 bp WT-GCDH band and/or a 390 bp KO-GCDH band.

### Cell culture

HEK293T were obtained from Dr. Eric Verdin (Gladstone Institute, San Francisco, CA) and maintained at 37 °C and 5% CO_2_ in complete medium: Dulbecco’s modified Eagle’s medium (DMEM, Gibco #11965118) supplemented with 10% (vol/vol) fetal bovine serum (FBS, Thermo Fisher Scientific #26140079). These cells were used to generate the HEK293T crSIRT5KO cell line with Dharmacon’s Edit-R gene engineering system (now part of Horizon Discovery) that uses plasmid-driven Cas9 nuclease expression, synthetic trans-activating CRISPR RNA (tracrRNA) and CRISPR RNA (crRNA) for gene of interest. As per manufacturer’s protocol, HEK293T cells at passage 17 were plated in 6-well plates at a density of 300,000 cells per well and were co-transfected with Edit-R hCMV-Puro-Cas9 (Dharmacon #U-005100-120), Edit-R tracrRNA (Dharmacon #U-002000-120) and Edit-R crRNA’s for SIRT5 using the DharmaFECT Duo (Dharmacon #T-2010-03) transfection reagent prepared in serum-free DMEM. After 48 h of transfection, cells were transferred to new 6-well plates and incubated in complete media containing 2 µg/mL puromycin for 3 days to positively select for transfected cells. Single cell monoclonal populations were generated from puromycin-selected cells using the 96-well serial dilution technique. Guide crRNA sequences (Table S1) targeting all known transcripts of SIRT5 (NM_012241_Exon5 310 aa; NM_031244_Exon5, 299 aa; NM_001193267_Exon5, 292 aa; and NM_001242827_Exon4, 202 aa) in the sense/anti-sense strands of human SIRT5 were selected from Dharmacon’s online CRISPR RNA configurator tool (http://dharmacon.gelifesciences.com/gene-editing/crispr-rna-configurator/: now https://dharmacon.horizondiscovery.com/gene-editing/crispr-cas9/crispr-design-tool/). Control cells underwent the same procedure as the SIRT5KO cells, except they did not receive the gene-specific crRNA. Each clonal population was cultured and screened for loss of SIRT5 protein by Western Blot. Ultimately, the confirmed SIRT5 crKO cell line was made using the crRNA ID ‘E’ (sequence in Table S1).

### Transfection and treatments

Plasmid DNA pcDNA3.1+ vector was obtained from Invitrogen (#V79020), pSG5 vector was from Agilent (#216201), WT-hSIRT5-HA was from Dr. Eric Verdin (Gladstone Institute, San Francisco, CA), and hGCDH-FLAG was cloned in-house in the pSG5 vector (as described below). GCDH-FLAG was overexpressed in HEK293T cells or the SIRT5 crWT/crKO cells over a period of 4 days (2 days of cell seeding followed by transfection and overexpression for 2 days). Cells are seeded in 100 mm dishes at 3 × 10^5 - 5 × 10^5 cells per dish and allowed to grow for in the complete medium for a period of 2 days. On day 2, cells growing in complete medium were transfected with a 1:1 mixture of plasmid DNA (7.5-15 µg) and Lipofectamine 2000 reagent (Invitrogen # 11668019); both prepared in OPTI-MEM (Gibco # 31985070) as per manufacturer’s recommendations. Plasmid DNA was allowed to overexpress for additional 2 days without any additional media change. For starvation conditions, cells on day 4 were washed 1X in PBS and either complete medium (control) or acylation medium (DMEM, no glucose, no glutamine, no pyruvate, no phenol red, no serum: Gibco # A1443001); or glutarylation media (Earle’s balanced salt solution (EBSS): Gibco # 24010043 + 5 mM glucose, + 50mM HEPES, + +0.8mM Lysine) was added. After 3 h incubation with complete, acylation, or glutarylation media, cells were either used for metabolic assays or washed 1x in PBS and collected in IP lysis buffer for immuno-precipitation experiments.

### Cloning

Human GCDH cDNA (clone ID 3142978) in pOTB7 vector was obtained from Thermo Fisher Scientific Open Biosystems repository (now part of Horizon Discovery). To clone into pSG5 vector, an EcoRI (enzyme, NEB # 0101S) site was introduced at the 5’ end using a forward primer: 5’-CAA AAA GAA TTC ATG GCC CTG AGA GGC GTC TC-3’. At the 3’ end, the original STOP codon was removed and a FLAG sequence followed by a new STOP codon was introduced. A BglII (enzyme, NEB # R0144S) restriction site was added at the 3’ end for cloning it into the pSG5 vector using a reverse primer: 5’-CAA AAA GAT CTT TAC TTG TCG TCA TCG TCT TTG TAG TCC TTG CTG GCC GTG AAC GCC TGG-3’. The PCR product was confirmed by EcoRI-BglII double restriction digestion and sequence verified using Sanger sequencing.

### Immuno-purification

Cells were lysed in IP lysis buffer (20 mM Tris base, 150 mM sodium chloride, 1 mM EDTA, 1 mM EGTA, 1% Triton-X 100) containing 10 mM nicotinamide, 10 mM sodium butyrate, 10 μM Trichostatin A (each 10 mL containing 100 μL each of phosphatase inhibitor cocktail 2 (Millipore-Sigma # P5726) and 3 (Millipore-Sigma # P0044), and 100 μL protease inhibitor cocktail (Millipore-Sigma # P8340) and the cell membranes were disrupted using a Series 60 Sonic Dismembrator (Model F60, Fisher Scientific, East Lyme, CT) with ten 2 s intermittent pulses. Soluble proteins were separated from insoluble cell debris by vortex mixing and centrifugation for 1 h at 4 °C and 14,000 × g. For overexpressed proteins, immuno-precipitation was performed using anti-FLAG-M2 resin (Millipore-Sigma # A2220) and 1-8 mg total soluble protein per IP reaction. Briefly, the resin was washed 3X in 1X TBS (tris-buffered saline) followed by 1X wash and re-suspention in lysis buffer. Soluble proteins in a total volume of 500-700 μL were incubated with pre-washed resin in IP spin columns (Pierce cat# 69725) overnight with constant rotation at 4 °C. Next day, the spin columns were spun at 10,000 × g for 2 min and the resin was further washed with IP lysis buffer 4X by inverting the tube for 10 times/wash. Resin-bound protein(s) were eluted by incubating samples with 50-100 μL of 2x Lane Marker non-reducing sample buffer (Thermo Fisher Scientific # 39001) at 95 °C for 5 min with open caps. Eluate was collected by spinning at 10,000 × g for 2 min, mixed with 2-mercaptoethanol (7.4% final concentration, Millipore-Sigma # M7522) and samples were re-heated at 95 °C for 5 min. Samples were run on 10% Tris-glycine SDS-PAGE gels (BIO-RAD #4561034) for maximum separation of proteins between 50 and 25 kDa. For immunoprecipitation of endogenous SIRT5 (Millipore-Sigma #HPA022002), GCDH (Millipore-Sigma # HPA048492), glutaryl-lysine- (Cell Signaling Technologies, non-commercial #14943MF), succinyl-lysine- (Cell Signaling Technologies, non-commercial # 13599) proteins, 1-8 mg of pre-cleared lysate/500 μL of lysis buffer was incubated with 2-10 μg of IP antibody overnight at 4 °C. About 20 μL of protein A/G beads (Thermo Fisher Scientific # 20423) were incubated with the antigen-antibody complex at room temperature for 4 h. Normal IgG was used as negative control in parallel with the IP reaction. For detection of endogenous immunopurified proteins, the primary antibody type (rabbit) was switched to mouse by using a conformation-specific mouse anti-rabbit antibody (Cell Signaling Technologies #3678) that only recognizes non-denatured antibodies to reduce the background from the heavy and light chains of the denatured IP antibody. The bands were then visualized by incubating with IRDye 800CW donkey anti-mouse IgG secondary antibody (LI-COR # 926-32212).

### Immunoblotting

Equal amounts of denatured protein samples were uniformly loaded and run on BioRad MiniPROTEAN or Criterion TGX Precast Midi Protein Gels, at 100-180 V for 45-60 minutes. Proteins were wet transferred to a 0.45 μm nitrocellulouse or Immobilon-P (Thermo Fisher Scientific, #88520) membranes in the Bio-Rad MiniPROTEAN or Criterion™ Blotter. Membranes were blocked for 60 minutes in LI-COR blocking buffer (1X TBS, 0.45% fish gelatin, 0.1% casein and 0.02% azide). Primary antibodies (1:1000-1:4000) were diluted in LI-COR blocking buffer containing 0.1-0.2% Tween-20 overnight at 4 °C. Next day, membranes were washed four-six times for 5-10 minutes each in 0.1% Tween-20 containing TBS. Infra-red dye-conjugated secondary antibodies were diluted 1: 10,000 in LI-COR blocking buffer containing 0.1-0.2% Tween-20 and incubated for 1 hour at room temperature. Western blots were visualized on an Odyssey CLx imager (LI-COR, Lincoln, NE). Un-stripped blots were re-probed for loading control and, image analysis for quantifying band-intensities was performed using the LI-COR Image Studio software (version 3.1). Antibodies: anti-MG-lysine (Millipore-Sigma # ABS2120-25UG), anti-SIRT5 (Millipore-Sigma #HPA022002), anti-GCDH (Millipore-Sigma # HPA048492), mouse anti-β-actin (Cell Signaling Technologies, #3700), rabbit anti-β-actin (Cell Signaling Technologies, #8457), anti-glutaryl-lysine (Cell Signaling Technologies, non-commercial #14943MF and PTM-Biolabs #1151), anti-succinyl-lysine (Cell Signaling Technologies, non-commercial # 13599 and PTM-Biolabs #401), anti-malonyl-lysine (Cell Signaling Technologies, #14942), anti-acetyl-lysine (Cell Signaling Technologies, #9441), IRDye 680RD donkey anti-mouse IgG (LI-COR # 926-68072), and IRDye 800CW donkey anti-rabbit IgG (LI-COR # 926-32213).

### Quantitative polymerase chain reaction (qPCR)

Total RNA was isolated from cells using the Qiagen’s RNeasy kit (# 74104) according to the manufacturer’s instructions (Qiagen, Germantown, MD). Total RNA (1 ug) was converted to cDNA using the iScript^TM^ cDNA synthesis kit (# 1708891, BIO-RAD, Hercules, CA) following the manufacturer’s protocol (BIO-RAD, Hercules, CA). Quantitative real-time PCR was performed on an Applied Biosciences ViiA^TM^ 8 thermocycler (Thermo Fisher Scientific) using iTaq™ Universal SYBR® Green Supermix (#1725121, BIO-RAD, Hercules, CA) and the following primer sets: SIRT3-Forward (5’-GCT GTA CCC TGG AAA CTA CAA-3’), SIRT3-Reverse (5’-ATC GAT GTT CTG CGT GTA GAG-3’), SIRT4-Forward (5’-GAG CTT TGC GTT GAC TTT CAG-3’), SIRT4-Reverse (5’-GGA CTT GCT GGC ACA AAT AAC-3’), SIRT5-Forward (5’-GCT CGG CCA AGT TCA AGT AT-3’), SIRT5-Reverse (5’-AAG GTC GGA ACA CCA CTT TC-3’), ATPSB-Forward (5’-CTA GAC TCC ACC TCT CGT ATC A-3’), and ATPSB-Reverse (5’-CAT ACC CAG GAT GGC AAT GA-3’). Relative amounts of SIRT3, SIRT4 and SIRT5 mRNA were calculated using the ΔΔCt method with ATP synthase subunit B as the endogenous control. Statistical comparisons were made using multiple t-test with Benjamini-Hochberg correction for multiple comparisons.

### Lysine oxidation assay

SIRT5WT and SIRT5KO cells were plated at 1*10^5 cells/well in 12-well plates in complete medium (DMEM/10% FBS). After 24 h, cells were washed 1X in PBS and 1 mL assay media containing 4μCi/mL U-[^14^C]-lysine (Moravek Inc #MC197 250UCI), 0.8 mM ^12^C-lysine, 50 mM HEPES, pH 7.5 in EBSS was added. Cells were incubated in this medium for 3 h after which 750 μL of the assay medium was aspirated and transferred to 13 mm culture tubes that contained a 200 µL 1N NaOH trap. The [^14^C]-CO_2_ generated as a result of lysine oxidation was released from the bicarbonate buffer by acidifying the medium with 70% perchloric acid. Tubes were incubated on a shaking water bath for 1 h at 37 °C. After 1 h, NaOH from the trap was carefully removed and added directly into scintillation vials containing the Uniscint BD scintillation fluid (National Diagnostics #LS-276). Disintegrations per minute were counted on a LS 6500 scintillation counter (Beckman Coulter, Fullerton, CA). The amount of label released was background subtracted and expressed as percentage of total radiolabel added in the incubation medium. Statistical analysis was performed using a two-tailed unpaired t-test to compare SIRT crWT vs SIRT crKO cells.

### Isolation of enriched mitochondrial fraction

Fresh mouse liver or cells were rinsed in ice-cold PBS and homogenized in 10 volumes of STE buffer (0.25 M sucrose, 10 mM Tris-HCl, pH 8.0, 1 mM EDTA, and 1X HALT protease inhibitor cocktail; Thermo Fisher Scientific #78420B) with 15 strokes in a chilled glass-teflon homogenizer. The homogenate was centrifuged at 700 × g for 10 min at 4 °C and the resulting supernatant centrifuged at 7,000 × g for 10 min at 4 °C. The pellet from the 7,000 × g centrifugation was used as the mitochondrial enriched fraction for glutaryl-CoA oxidation assay and blue-native PAGE.

### Glutaryl-CoA oxidation assay

Mitochondria were isolated from fresh livers of male SIRT5WT and SIRT5KO mice (3-4 month old with GCDHKO mouse liver mitochondria as negative control) or cells as described above. Mitochondria were permeabilized by resuspending the mitochondrial pellet in permeabilization buffer (105 mM K-MES, pH 7.1, 30 mM KCl, 10 mM KH_2_PO_4_, 5 mM MgCl_2_, 0.5 mg/ml BSA, and 30 µg/ml alamethicin; Sigma-Aldrich #A5362) at a protein concentration of 3-5 µg/µL and incubated for 5 min on ice. GCDH activity assay was initiated by mixing 100 ul of permeabilized mitochondria with 100 µl of GCDH reaction buffer (105 mM K-MES, pH 7.1, 30 mM KCl, 10 mM KH_2_PO_4_, 5 mM MgCl_2_, 0.5 mg/ml BSA, 30 µg/ml alamethicin, 200 µM FAD, and 0.1 µCi/µl [1, 5-^14^C]-glutaryl-CoA; American Radiolabeled Chemicals Inc. # ARC3471). The reaction was incubated for 30 min at 37 °C and the [^14^C]-CO_2_ produced was trapped and quantified as described for the lysine oxidation assay. Statistical analysis was performed using a two-tailed unpaired t-test to compare SIRTWT Vs SIRTKO mouse liver lysates.

### Blue native polyacrylamide gel electrophoresis (BN-PAGE)

Liver mitochondria were isolated from 24-h fasted male SIRT5WT and SIRT5KO mice (2 month old, age matched GCDHWT and GCDHKO mice were used as controls) using the above-mentioned procedure and stored at −80 °C. Thawed mitochondrial pellets were solubilized in 1X Native-PAGE sample buffer containing 2% n-dodecyl-β-D-maltoside, 0.5% G-250, and 1X inhibitors (protease inhibitors, phosphatase inhibitors, nitcotinamide, trichostatin A and sodium butyrate) as per manufacturer guidelines for NativePAGE™ Sample Prep Kit (ThermoFischer Scientific #BN2008). Equal amounts of protein (15 µg) were loaded on NativePAGE™ Novex® 4–16% Bis-Tris Gels and were run at 150 V for first 60 minutes followed by another 60 min at 250 V. Proteins were transferred to PVDF membranes by wet transfer method mentioned above. Post-transfer PVDF membranes were washed 3X in 100% methanol for 5 min each to remove excess G250 dye and were processed for immunoblotting as mentioned above. Statistical analysis was performed using a two-tailed unpaired t-test to compare SIRTWT Vs SIRTKO mouse liver lysates.

### Chemical acylation and deacylation of recombinant human GCDH protein

Recombinant human GCDH was expressed and purified to homogeneity as described in (18). Glutaryl-GCDH (or succinyl-GCDH) was prepared by incubating the protein with excess glutaryl-CoA (or succinyl-CoA). Briefly, 20 μM GCDH was incubated with 1.6 mM glutaryl-CoA/succinyl-CoA, in acylation buffer (50 mM HEPES pH 8.0 and 150 mM NaCl) for 15 h at 25 °C, with gentle stirring. After incubation, the reaction mixture was applied to a Superdex 200 Increase 10/300 GL gel filtration column (GE Healthcare Life Sciences #28-9909-44) to remove excess glutaryl-CoA/succinyl-CoA. Modified GCDH was eluted at a rate of 0.75 ml/min with elution buffer (25 mM HEPES pH 7.8 and 30 mM NaCl). The Superdex 200 column was previously calibrated using standard proteins (Ferritin, ovalbumine, ribonuclease A, conalbumin and carbonic anhydrase) which were used to determine the molecular mass of the pure modified proteins. The tetrameric form of the pure unmodified/modified proteins was used in subsequent structural and enzymatic activity assays. For deacylation recombinant 22.5 μg SIRT5, (9) was incubated with glutaryl-GCDH or succinyl-GCDH (135 μg), in 5 mM Tris-HCl pH 9, 4 mM MgCl_2_, 5 mM NaCl, 0.5 mM DTT, 1 mM NAD^+^, for 3 h at 37 °C. Unmodified GCDH and modified-GCDH with or without SIRT5 were used as controls for this deacylation reaction.

### GCDH activity assay

GCDH activity was evaluated using two distinct enzymatic assays. In the first assay, GCDH activity was measured at 25 °C, monitoring the reduction of 2,6-dichlorophenolindophenol (DCPIP, 30 μM, Millipore-Sigma # 36180) at 600 nm, upon energization with the substrate glutaryl-CoA (15 μM) in the presence of the phenazine methosulfate (PMS, 2 mM, Millipore-Sigma # P9625) mediator, as in (18). In the second assay, the activity of GCDH was inferred from its ability to reduce the electron transfer flavoprotein (ETF), which is its physiological electron acceptor. For this, GCDH was incubated with glutaryl-CoA (15 μM) and purified recombinant human ETF (19, 20) and the ability of ETF to receive electrons from GCDH was monitored by measuring DCPIP reduction at 600 nm (21). Buffer for enzymatic assays was 10 mM HEPES pH 7.8. One unit of catalytic activity is defined as nmol of DCPIP reduced per minute, under conditions used in the assay. The activity of glutarylated- and deglutarylated-GCDH was calculated as percentage of unmodified GCDH controls. All measurements were made in quadruplicates and statistical analysis was performed using a one way ANOVA with Tukey’s post hoc test to make multiple comparisons between unmodified, modified and de-acylated (SIRT5 co-incubation) GCDH samples.

### Spectroscopic methods

Fluorescence spectroscopy was performed using a Jasco FP-8200 spectrofluorometer with a cell holder thermostatically controlled with a Peltier. For tryptophan emission excitation wavelength was set at 280 nm, and FAD emission was followed by setting excitation wavelength at 450 nm; slits were 5 and 10 nm for excitation and emission, respectively. Typically, protein concentration was 2 μM. CD spectra were recorded for GCDH on a Jasco J-1500 spectropolarimeter with a cell holder thermostatically controlled with a Peltier. A quartz polarized 1 mm path length quartz cuvette (Hellma) was used, and protein concentrations were typically 0.1 mg/ml.

### Thermal stability

Thermal unfolding with a linear temperature increase was followed using circular dichroism (ellipticity variation at 222 nm) and fluorescence spectroscopy (tryptophan emission λ_ex_ =280 nm and λ_em_=340 nm, and FAD emission λ_ex_ =450 nm and λ_em_ =530 nm). In all experiments, a heating rate of 1 °C/min was used, and temperature was increased from 20 to 85 °C. Data were analyzed according to a two-state model, and fits to the transition curves were made using OriginPro8.

### Label-free qualitative acyl-proteomic analysis of recombinant GCDH

Chemically glutarylated and succinylated human recombinant GCDH protein was prepared as mentioned above. About 10-30 ug protein was loaded on 10% NuPAGE Bis-Tris protein gels (Thermo Fischer Scientific #NP0301BOX) and separated using the NuPAGE™ MOPS SDS running buffer (Thermo Fischer Scientific #NP0001) for 1 h at 180 volts. Post-run, the gel was stained overnight in colloidal blue stain (Thermo Fischer Scientific #LC6025), washed in de-ionized water, and bands corresponding to ~44 kDa were excised and collected in 1.5 mL micro-centrifuge tubes. Gel pieces were cut into smaller pieces using a sterile 20 µL pipette tip, vortex mixed and incubated twice in 200 µL of a 1:1 mixture of acetonitrile (ACN) and 100 mM ammonium bicarbonate (AmBIC) at room temperature for 15 min followed by dehydration in 200 µL of 100% ACN from 1min. The gel pieces were rehydrated and reduced with 200 µL 10 mM dithiothreitol (DTT) in 100 mM AmBIC for 30 min at 55 °C. After discarding the DTT-solution, the samples were alkylated using 200 µL of 55 mM iodoacetamide in 100 mM AmBIC in dark at room temperature for 30 min. Samples were further washed in 200 µL of 1:1 ACN: 100 mM AmBIC mixture followed by 200 uL of 100% ACN. Samples were rehydrated in 200 µL of digestion buffer (50 mM AmBIC, 5 mM CaCl2) containing 1 ug Trypsin per 50 ug of GCDH protein and in-gel digested overnight at 37 °C. Trypsin digested peptides were serially extracted using 200 µL each of 50% ACN, 0.3% formic acid and 80% ACN, 0.3% formic acid at room temperature for 15 min each. Pooled extracts were subjected to speed-vac until dry and resuspended in 1 mL of 0.5% trifluroacetic acid. Acidified samples were then desalted by solid phase extraction using 50 mg tC18 Sep-Pak columns (Waters) and dried using speed-vac.

For assessment of glutarylated and deglutarylated GCDH, samples were resuspended in 22 µL of 0.1% formic acid and subjected to *nano*LC-MS/MS analysis using a using an EASY-nLC UPLC system (Thermo Fisher Scientific) coupled to a *Q Exactive Plus* Hybrid Quadrupole-Orbitrap mass spectrometer (Thermo Fischer Scientific) *via* a nanoelectrospray ionization source. To minimize effects of acylpeptide carryover, we assessed unmodified GCDH first and blanks in between samples. Sample injections of 10 µL were first trapped on an Acclaim PepMap 100 C18 trapping column (3 um particle size, 75 µm × 20 mm) with 22 µL of solvent A (0.1 % FA) at a variable flow rate dictated by max pressure of 500 Bar, after which the analytical separation was performed over a 105 minute gradient (flow rate of 300 nL/minute) of 5 to 40% solvent B (90% ACN, 0.1% FA) using an Acclaim PepMap RSLC C18 analytical column (2 um particle size, 75 µm × 500 mm column (Thermo Fischer Scientific) with a column temperature of 55°C. MS^1^ (precursor ions) was performed at 70,000 resolution, with an AGC target of 3 × 10^6^ ions and a maximum injection time (IT) of 100 ms. MS^2^ spectra (product ions) were collected by data-dependent acquisition (DDA) of the top 20 most abundant precursor ions with a charge greater than 1 per MS1 scan, with dynamic exclusion enabled for a window of 30 seconds. Precursor ions were filtered with a 1.2 m/*z* isolation window and fragmented with a normalized collision energy (NCE) of 27. MS2 scans were performed at 17,500 resolution, with an AGC target of 1×10^5^ ions and a maximum IT of 100 ms. Similar methods were used for direct comparison of PTMs resulting from glutaryl and succinyl-CoA reactions.

Raw LC-MS/MS data were processed in Proteome Discoverer v2.2 (PD2.4, Thermo Fisher Scientific), using the Sequest HT and MS Amanda 2.0 (Protein Chemistry Facility IMP/IMBA/GMI, Vienna, Austria.) search engines as nodes. Data were searched against the UniProt *E. Coli* (strain K12) complete proteome database of reviewed (Swiss-Prot) and unreviewed (TrEMBL) proteins, which consisted of 4,276 sequences on the date of download (3/15/18) and hGCDH-6XHIS (modified from UniProtKB - Q92947). Variable modifications included oxidation of methionine (M) and glutarylation (114.0316941 Da monoisotopic mass) of lysine (K), with carbamidomethyl of cysteine (C) as a fixed modification. Data were searched with a 10 ppm precursor mass and 0.02 Da product ion tolerance. The maximum number of missed cleavages was set at 4 and enzyme specificity was designated as trypsin. Peptide spectral matches (PSMs) were filtered to a 1% false discovery rate (FDR) with Percolator (22). Site localization probabilities were determined using ptmRS (23) using a 75% Site Probability Threshold. PSMs were grouped to peptides maintaining 1% FDR at the peptide level and peptides were grouped to proteins using the rules of strict parsimony. Proteins were filtered to 1% FDR using Protein FDR Validator. Peptide quantification was done using the MS1 precursor intensity from aligned features (10 min max retention time shift) and imputation was performed via low abundance resampling. Quantitation for each GCDH acylpeptide identified was normalized to the relative abundance of hGCDH-6XHIS protein (calculated from unmodified GCDH peptides only) to control for any slight deviations in sample loading and LC-MS performance. Statistical significance was assessed by taking the Log2 of GCDH-normalized acylpeptide abundance values, and calculating Log2 fold-changes and performing the student’s t-test (n=3) across conditions. The mass spectrometry proteomics data have been deposited to the ProteomeXchange Consortium via the PRIDE (24) partner repository with the dataset identifier PXD018156.

### Molecular modeling of GCDH-acyl sites

The crystal structure coordinates for the homotetramer of human glutaryl-CoA dehydrogenase (25) were downloaded from the Protein Data Bank (PDB ID: 1SIR; www.rcsb.org; validation report: https://files.rcsb.org/pub/pdb/validation_reports/si/1sir/1sir_full_validation.pdf). The structure was assessed to correct the protonation states, predict side chain pKa, optimize the intra-/ inter-molecular hydrogen bonding networks (26, 27), and add the two missing C-terminal residues to each subunit using YASARA Structure v17.4 (www.yasara.org). The protein structure was then typed with the CHARMM force field (28) and subjected to energy minimization with the Generalized Born with simple switching (GBSW) implicit-solvent model to a root mean square (RMS) convergence of < 0.01 kcal/mol using Biovia Discovery Studio 2018 (www.3dsbiovia.com). Figures were generated using Lightwave3D 2018 (www.lightwave3d.com).

### Targeted Metabolomics

293T SIRT5crWT and crKO cells were collected during the exponential growth phase and metabolites were isolated and analyzed using stable isotope dilution technique. Amino acids and acylcarnitine measurements were made by flow injection tandem mass spectrometry using sample preparation methods described previously (29, 30). The data were acquired using a Waters TQD mass spectrometer equipped with Acquity^TM^ UPLC system and controlled by MassLynx 4.1 operating system (Waters, Milford, MA).

### Non-targeted Metabolomics in SIRT5 CRISPR cells

Non targeted metabolomics were performed using Sirt5 crWT and crKO 293T cells that were either fed (complete DMEM, (Gibco # 11965118) supplemented with 10% (vol/vol) fetal bovine serum (FBS, Thermo Fisher Scientific #26140079)), fasted (1 hour EBSS with phenol red) or refed (fasted 1 hour in EBSS with phenol red and refed for 1 hour in complete DMEM). Cells were prepared for metabolomics by first removing media and then placing on dry ice and adding pre-cooled 80% methanol/water (v/v; HPLC grade). Enzymes were further inactivated by incubating plate(s) in −80°C freezer 15 minutes. Cells were scraped into the solvent, transferred to tubes and centrifuged at 20,000 rcf, 10min at 4°C. Samples were dried using a speed vacuum at room temperature and pellets were stored at −80°C until processing for LC-MS.

Metabolite extraction was performed as described in previous study (31). The supernatant was transferred to a new Eppendorf tube and dried in vacuum concentrator at room temperature. The dry pellets were reconstituted into 30 ml sample solvent (water:methanol:acetonitrile, 2:1:1, v/v) and 3 ml was further analyzed by liquid chromatography-mass spectrometry (LC-MS). Ultimate 3000 UHPLC (Dionex) is coupled to Q Exactive Plus-Mass spectrometer (QE-MS, Thermo Scientific) for metabolite profiling. A hydrophilic interaction chromatography method (HILIC) employing an Xbridge amide column (100 × 2.1 mm i.d., 3.5 mm; Waters) is used for polar metabolite separation. Detailed LC method was described previously(32) except that mobile phase A was replaced with water containing 5 mM ammonium acetate (pH 6.8). The QE-MS is equipped with a HESI probe with related parameters set as below: heater temperature, 120 °C; sheath gas, 30; auxiliary gas, 10; sweep gas, 3; spray voltage, 3.0 kV for the positive mode and 2.5 kV for the negative mode; capillary temperature, 320 °C; S-lens, 55; A scan range (m/z) of 70 to 900 was used in positive mode from 1.31 to 12.5 minutes. For negative mode, a scan range of 70 to 900 was used from 1.31 to 6.6 minutes and then 100 to 1,000 from 6.61 to 12.5 minutes; resolution: 70000; automated gain control (AGC), 3 × 106 ions. Customized mass calibration was performed before data acquisition. LC-MS peak extraction and integration were performed using commercial available software Sieve 2.2 (Thermo Scientific). The peak area was used to represent the relative abundance of each metabolite in different samples. The missing values were handled as described in previous study (32).

### Bioinformatics

The code for gene correlation is available within the Hirschey Lab github (https://github.com/matthewhirschey/livercoexpression).To summarize, liver gene expression values, IDs, and descriptions were downloaded from [NCBI GEO]; (https://www.ncbi.nlm.nih.gov/geo/query/acc.cgi?acc=GSE14520). Data were cleaned, tidied, and prepared for analysis within R using the ‘tidyverse’ package. Liver samples were filtered to include only non-tumor samples. A correlation matrix was built to identify similar patterns of hepatic expression for all genes compared to all genes using the ‘corrr’ package. Probe IDs for SIRT3, 4, and 5 were queried and summarized for the top 20 gene correlations. To measure the distribution of correlation coefficients, r^2^ values were resampled from the entire matrix to generate 1000 ‘virtual’ gene lists using the package ‘moderndive’. Mean and standard deviations were calculated from this virtual set. A standard deviation threshold of +/− two standard deviations was set, and the resulting gene correlations with each gene of interest greater than or less than the threshold were considered to be statistically unlikely by chance, and then queried for gene set enrichment analysis.

### Statistical analyses

Unless otherwise indicated, figure data represent the mean ± standard deviation. Significance for comparison between two groups was evaluated using a two-tailed unpaired Student’s t-test or a one-way ANOVA followed by appropriate post-hoc test (parametric or non-parametric) for comparisons between more than two groups. A result was considered significant if p ≤ 0.05. Statistical tests and graphs were prepared using GraphPad Prism 7.0 software

### Data Availability

Proteomic data files have been deposited on ProteomeXchange with the accession number PXD018156. Code for computational analyses is available on github (https://github.com/matthewhirschey/livercoexpression).

## ACKNOWLEDGEMENTS

We would like to acknowledge members of the Hirschey laboratory for discussion and comments, and Zhihong Lin for assistance with animal husbandry; James Draper for his assistance with proteomic data analysis; Cell Signaling Technology (Danvers, MA) for providing succinyl-, glutaryl-, and malonyl-lysine antibodies, valuable discussions and technical support; and Jason Locasale for assistance with non-targeted metabolomic experiments. We would like to acknowledge funding support from The Ellison Medical Foundation, the Glenn Foundation, the National Institutes of Health and the NIDDK grant R01DK115568, the National Institutes of Health and the NIA grant R01AG045351. Computational simulations were conducted at the University of Colorado Computational Chemistry and Biology Core Facility, which is funded in part by NIH/NCATS Colorado CTSI Grant UL1TR001082. The GCDHKO mouse strain used for this research project, B6.129S4-Gcdhtm1Dmk/Mmnc, identification number 034368-UNC, was obtained from the Mutant Mouse Regional Resource Center, a NIH funded strain repository (8U42OD010924-13), and was donated to the MMRRC by David M. Koeller, MD, Oregon Health & Science University. The content is solely the responsibility of the authors and does not necessarily represent the official views of the National Institutes of Health or other funding sources. This work was partly supported by the Fundação para a Ciência e Tecnologia (FCT/MCTES, Portugal) through research grant PTDC/BBB-BQB/5366/2014 (to B.J.H.), and fellowship SFRH/BPD/74475/2010 (to B.J.H.), and SFRH/BD/89690/2012 (to T.G.L.). The Gomes laboratory is partly supported by grant UIDB/ 04046/2020, research unit grant from FCT, Portugal (to BioISI).

## AUTHOR CONTRIBUTIONS

D.P.B., K.A., M.D.H. conceptualized the study. Investigation, D.P.B., C.A.M., K.A., B.J.H., T.G.L., S.F., H.G., D.S.B., M.P.S., P.A.G. A.A., S.K., J.L.; Writing - Original Draft, D.P.B. C.A.M and, M.D.H.; Writing-Review & Editing, All authors; Supervision, C.M.G., M.D.H.; Project Administration, M.D.H.; Funding Acquisition, B.J.H., C.M.G., and M.D.H.

## ADDITIONAL INFORMATION

Correspondence and requests for materials should be addressed to MDH

---

**Table S1:**
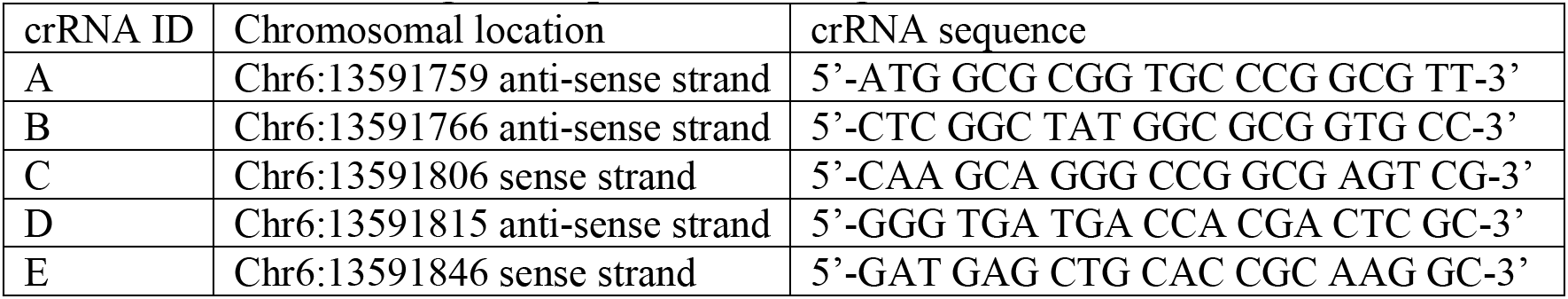
CRISPR RNA guide sequences and target chromosomal location.

**Table S2: List of acylation sites on Recombinant GCDH Related to Figure 1**

Supplementary Excel file containing label-free quantitative mass spectrometry data assessing glutarylation of lysines on hGCDH-6xHis after chemical glutarylation and incubation with or without mSirt5. Data are shown on the following spreadsheet tabs:

**Tab 1) GCDH Glutaryl Site Summary:** Summary of relative quantitation statistics for glutarylation at individual sites on hGCDH-6His, calculated from a subset of peptides identified at 1% FDR (Tab2). Some sites combine quantitation from multiple GCDH glutaryl-peptides, all normalized for hGCDH-6xHis abundance in each sample (calculated from unmodified GCDH peptides only).

**Tab 2) Peptide Groups:** Output from Proteome Discoverer 2.4, showing information on all peptides identified at 1% FDR. Column J highlights peptides mapping to hGCDH-6His (yellow) or mSirt5 (orange), while column Q highlights site localization probabilities for GCDH glutaryl-peptides (green). The “Abundances (Normalized)” values (columns AO-AW) were used for summing quantification for peptides with common glutaryl-sites, applying log_2_-normalization, and testing for statistical significance (Tab1).

**Tab 3) Protein Groups:** Output from Proteome Discoverer 2.4, showing information on all “Master Proteins”, or representatives of protein groups to which the peptides in Tab2 map at 1% protein-level FDR. Rows are colored to highlight hGCDH-6His (yellow) and mSirt5 (orange) and are rank ordered by Coverage (Column N), with hGCDH—6His as the top protein at 100% sequence coverage. The hGCDH-6His (Row 2, yellow) “Abundances (Normalized)” values (columns BI-BQ) are equal for all samples as all quantitation was normalized to hGCDH-6His levels in each sample.

**Table S3: crSIRT5 raw metabolomics data**

This table contains the raw metabolomics data associated with Figure 3. Analysis of these data is detailed in the Non-targeted metabolomics section of the materials and methods.

## SUPPORTING INFORMATION

**Supporting Figure S1:**
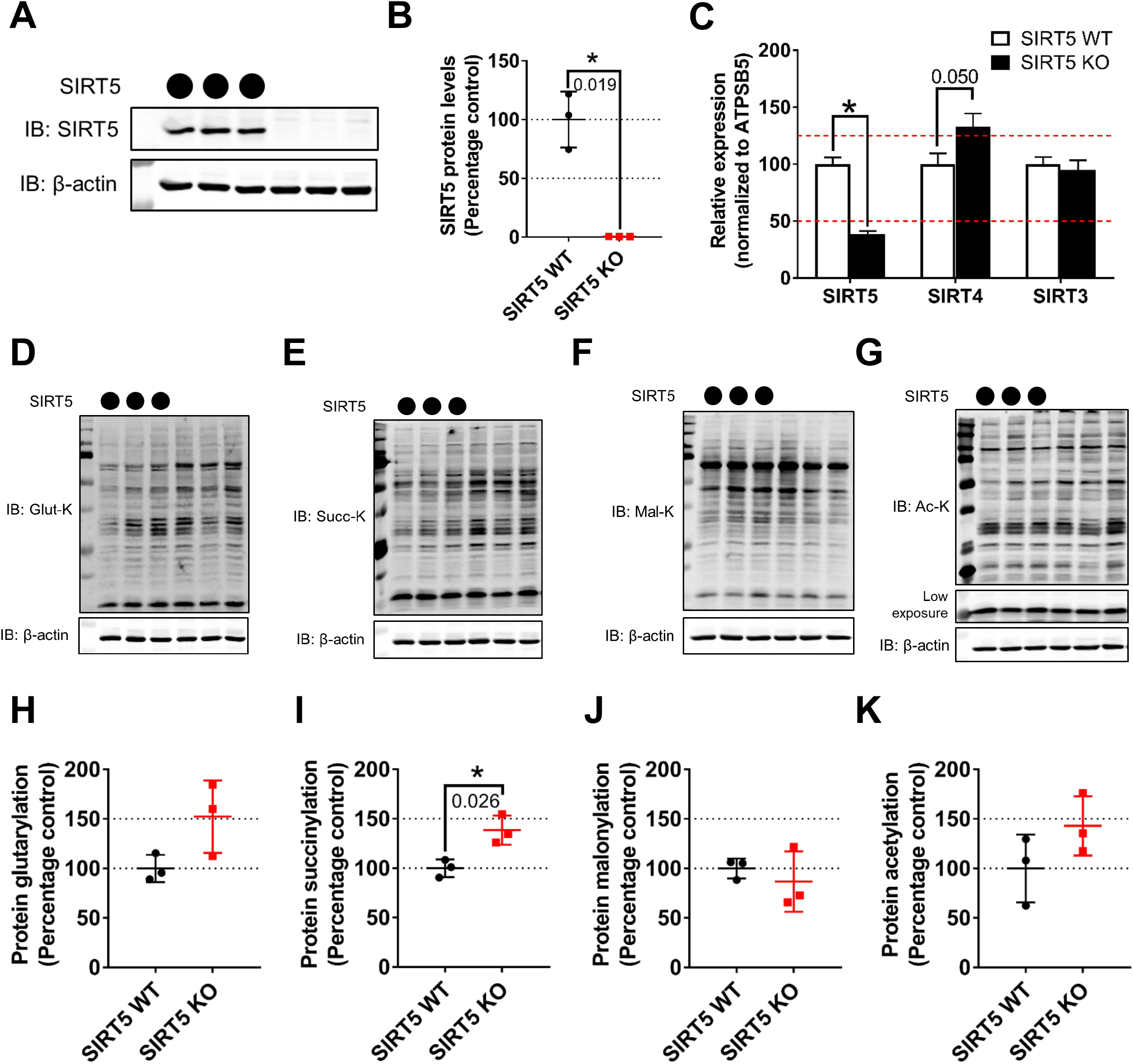
Validation of HEK293T CRISPR SIRT5 KO cells. (A) Immunoblot for SIRT5 protein from HEK293T crWT (circles) clone, and crKO (blank) clone (using CRISPR guide E, table S1). (B) Relative SIRT5 protein quantification of (A) (mean ± SD, n=3). (C) Relative mRNA expression of mitochondrial sirtuins (SIRT3, 4 and 5) normalized to ATP subunit 5 mRNA levels in 293T SIRT5 crWT and crKO cells (mean ± SEM, n=5, *= p-value ≤ 0.05). (D-G) Immunoblots of SIRT5 crWT and crKO whole cell lysates blotted using antibodies against glutaryl-lysine (D), -succinyl-lysine (E), malonyl-lysine (F), and acetyl-lysine (G). (H-K) Relative quantification of acylated proteins normalized to β-actin levels: glutarylation (H), succinylation (I), malonylation (J), and acetylation (K) (mean ± SD, n=3).

**Supporting Figure 2 (related to Figure 1):**
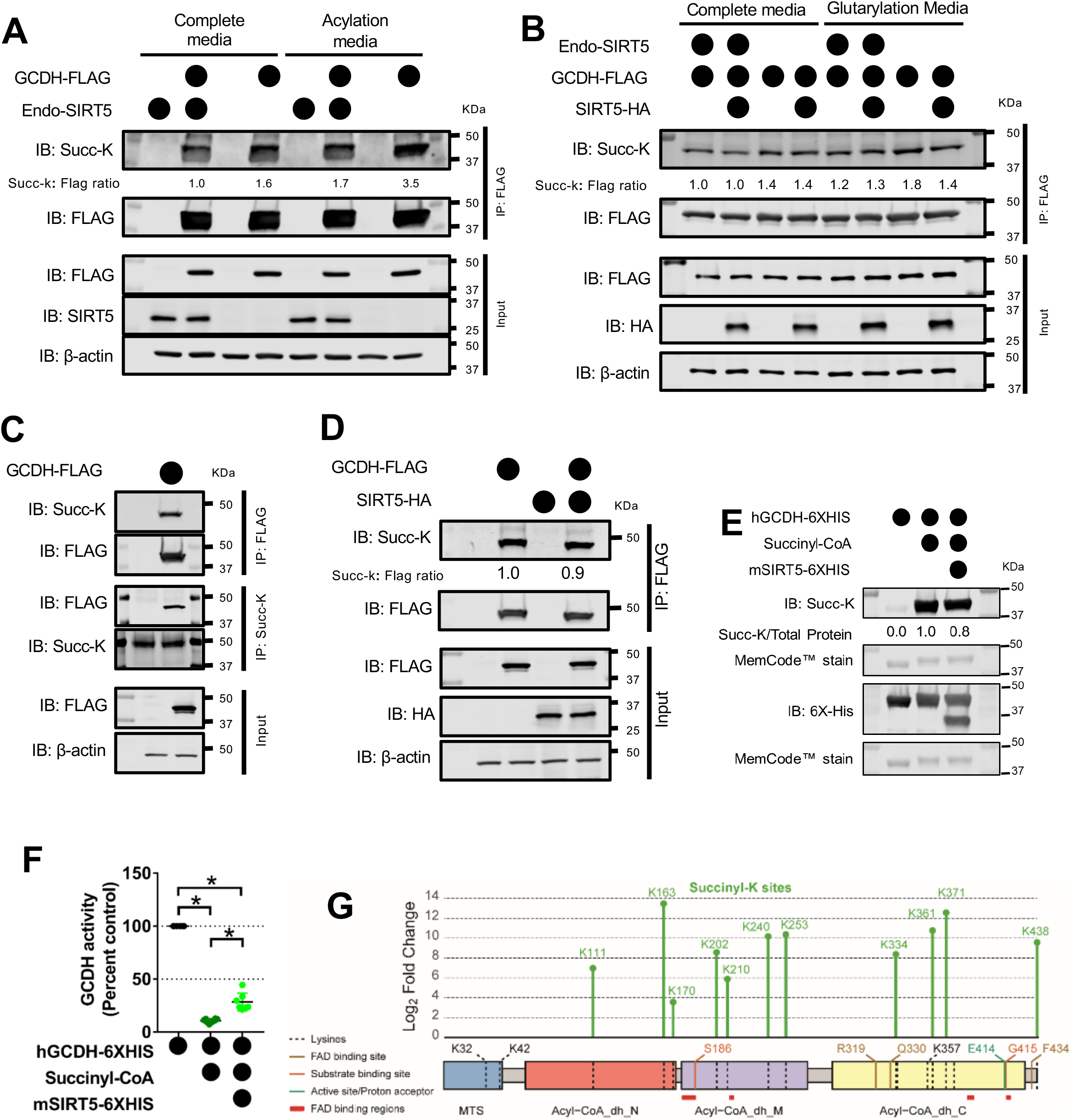
GCDH Succinylation. (A) Immunoblot for immuno-purified GCDH-FLAG by Flag-M2 resin from 293T SIRT5 crWT or crKO cells grown in complete media or acylation media (DMEM - glucose, - glutamine, - pyruvate, + 10% FBS). Blots are representative of at least three independent experiments and quantitative values of succinyl-lysine intensity normalized to FLAG intensity expressed as ratios relative to control (overexpressed GCDH-FLAG in SIRT5 crWT cells grown in complete media, lane 2. Same experiment as Figure 1C, blotted for Succ-K). (B) Immunoblot for immuno-purified GCDH-FLAG by Flag-M2 resin from SIRT5 crWT or crKO cells grown in complete media or glutarylation media (EBSS +50mM HEPES, +0.8mM Lysine) with or without co-expressed SIRT5-HA. Blots are representative of at least three independent experiments and quantitative values of glutaryl-lysine intensity normalized to FLAG intensity expressed as ratios relative to control (overexpressed GCDH-FLAG in SIRT5 crWT cells grown in complete media, lane 1. Same experiment as Figure 1D, blotted for Succ-K). (C) Immunoblot for immuno-purified GCDH-FLAG by Flag-M2 resin or succinyl-proteins by anti-succinyl-lysine antibody (Succ-K) from 293T SIRT5 CRISPR cells grown in complete media media. Blots representative of at least three independent experiments. (D) Immunoblot of immuno-purified GCDH-FLAG by Flag-M2 resin from 293T SIRT5 CRISPR cells grown in complete media with or without co-expressed SIRT5-HA. Blots representative of at least three independent experiments and quantitative values of succinyl-lysine intensity normalized to FLAG intensity expressed as rations relative to control (overexpressed GCDH-FLAG, lane 2. Same experiment as Figure 1B, blotted for Succ-K) (E) Immunoblot of chemically modified GCDH (hGCDH-6xHIS) using succinyl-CoA with or without incubation with recombinant SIRT5 (mSIRT5-6xHIS). Quantitative values of succinyl-lysine intensity normalized to total protein are expressed as ratios relative to control (succinyl-modified recombinant GCDH, lane 2) (F) Enzymatic activity of unmodified (black), succinylated (dark green) and desuccinylated (bright green) GCDH determined by using PMS/DCPIP as artificial electron acceptors and Succinyl-CoA as electron donor (mean ± SD, n=6, 8, and 7 for unmodified, modified and deacylated GCDH respectively; * = p-value ≤ 0.05). (G) Succinyl-K sites as identified by label-free quantitative LC-MS/MS on recombinant GCDH are mapped on full-length human GCDH protein.

**Supporting Figure 3:**
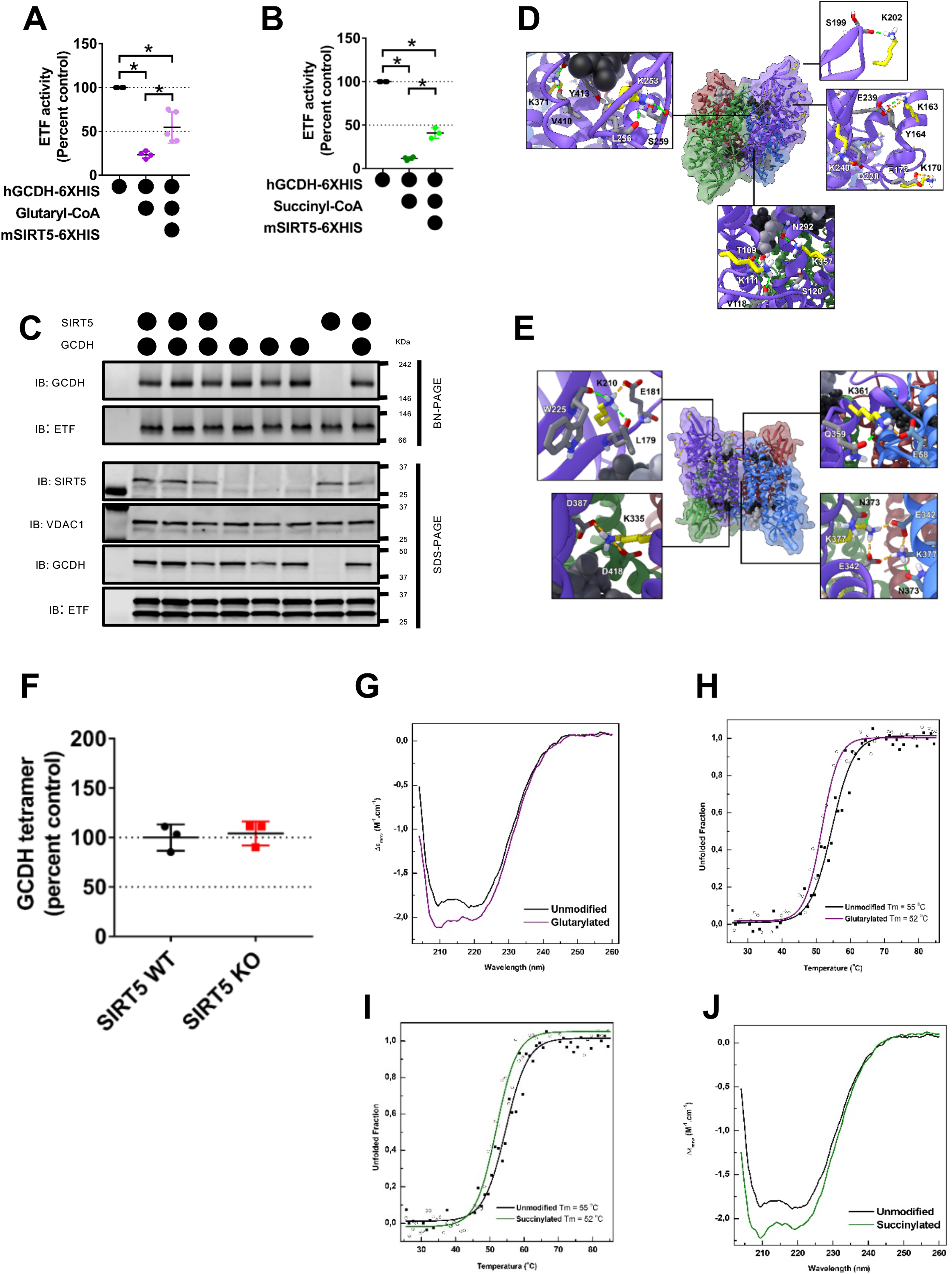
Acylation induces modest alterations in GCDH structure. (A) Enzymatic activity of unmodified (black), glutarylated (dark purple) and de-glutarylated (light purple) GCDH determined by using ETF/DCPIP as an electron acceptors and Glutaryl-CoA as electron donor (mean ± SD, n = 4, 4, and 5 for unmodified, modified and deacylated GCDH respectively, **\*:** p-value ≤ 0.05). (B) Enzymatic activity of unmodified (black), succinylated (dark green) and de-succinylated (light green) GCDH determined by using ETF/DCPIP as an electron acceptors and Succinyl-CoA as electron donor (mean ± SD, n = 4, 4, and 3 for unmodified, modified and deacylated GCDH respectively, **\*:** p-value ≤ 0.05). (C) Immunoblot of blue-native PAGE separated GCDH tetramer and SDS-PAGE for denatured GCDH isolated from liver mitochondria of 24 h fed SIRT5WT and SIRT5KO mice. Fed GCDHKO mouse liver mitochondrial lysates used as control to identify correct GCDH tetramer complex. (D-E) Homotetrameric structure of human GCDH with each subunit colored separately (purple, blue, red, and green). Yellow sticks indicate modified lysine residues. The FAD cofactor and CoA substrate are black and light gray spheres, respectively. (F) Quantitation of GCDH tetramer intensity normalized to ETF complex intensity. Data representative of two independent experiments (mean ± SD, n = 3 for both SIRT5 WT and SIRT5 KO). (G) Far-UV CD spectra for glutarylated-GCDH (purple) and unmodified GCDH (black). (H) Thermal stability profiles for glutarylated-GCDH (open circles, purple curve) and unmodified GCDH (closed squares, dark curve). The solid lines represent two-state sigmoid curves from which the apparent midpoint temperature were determined. (I) Far-UV CD spectra for succinylated-GCDH (green) and unmodified GCDH (black). (J) Thermal stability profiles for succinylated-GCDH (open circles, green curve) and unmodified GCDH (closed squares, green curve). The solid lines represent two-state sigmoid curves from which the apparent midpoint temperature was determined.

**Supporting Figure 4 (related to Figure 4):**
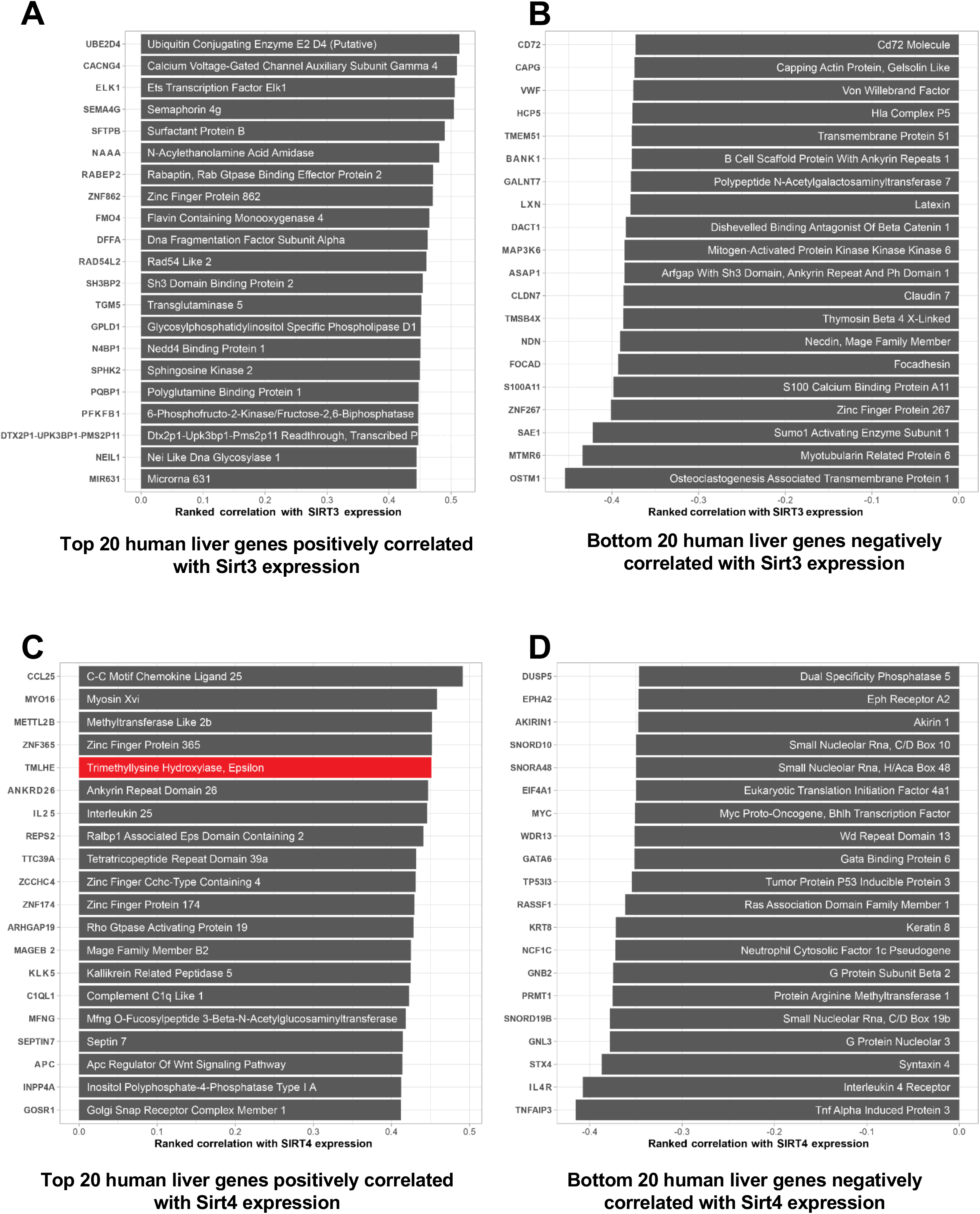
Mouse liver co-expression analysis for SIRT3 and 4. (A-B) Human liver Sirt3 liver co-expression analysis showing gene expressions that positively (A) or negatively (B) correlate to Sirt3 gene expression. (C-D) Human liver Sirt4 liver co-expression analysis showing gene expressions that positively (C) or negatively (D) correlate to Sirt4 gene expression. Genes highlighted in red have known roles in amino acid metabolism.

